# Complex adaptive architectures constrain the pace of adaptations sweeping across human gut microbiomes

**DOI:** 10.64898/2026.06.01.729358

**Authors:** Peter J. Laurin, Nandita R. Garud

## Abstract

Recent work has shown that commensal gut bacteria can evolve rapidly within hosts on short timescales of days to months, fueled by the enormous mutational input generated daily in the microbiome. Yet how rapidly adaptations spread across gut microbiomes of different hosts remains unclear. We address this question by estimating the number of independent origins of gene-specific sweeps spreading via recombination across bacterial populations. Multiple origins (soft sweeps) indicate that adaptive mutations arise rapidly whereas one origin (hard sweeps) indicate slower mutational input. Contrary to expectations of rapid adaptation, we find that many gene-specific sweeps have only one or a few origins. We show that this requires that sweeps arise from adaptive mutation rates orders of magnitude lower than single base pair mutation rates. This implies that gene-specific sweeps bear difficult-to-mutate complex adaptations such as structural or epistatic variants. Consistent with this interpretation, we find that identified sweep regions exhibit patterns of nucleotide diversity and linkage disequilibrium inconsistent with a single adaptive mutation rising to high frequency. We conclude that recombination across human gut microbiomes enables the spread of adaptations with complex genetic architectures that otherwise would require a long waiting time to generate *de novo* within an individual host.

## Introduction

Each human gut microbiome is estimated to experience billions of new mutations daily (Zhao et al. 2019). Consistent with this massive mutational input, recent work has shown that commensal gut bacteria can rapidly evolve within hosts on short timescales of days to months, even in healthy individuals without any obvious selection pressures, such as antibiotics (Garud et al. 2019; Zhao et al. 2019; Poyet et al. 2019; Yaffe and Relman 2019; Groussin et al. 2021; Ghalayini et al. 2018). However, far less is known about the rapidity of the spread of adaptations across human gut microbiomes, making it challenging to predict when and how a gut microbiome will evolve an adaptation.

An adaptive mutation that is beneficial in multiple hosts may arise *de novo* in distinct individuals’ microbiomes. However, an important complication is that *de novo* mutations may not be the only source of new genetic variation: resident strains may acquire adaptations via recombination with another strain already bearing the adaptive allele (Liu and Good 2024; Frazão et al. 2019). If broadly adaptive across multiple microbiomes, adaptive fragments may recombine onto multiple strain backgrounds, resulting in a ‘gene specific sweep,’ in which both the adaptive allele and linked variants in the nearby vicinity rise to high frequency. Recently, we found that such recombination-mediated gene-specific selective sweeps are pervasive and have swept across human gut microbiomes globally (Wolff and Garud 2026).

However, the extent of gene-specific sweeps is surprising: the extremely large input of *de novo* mutations should in theory allow the same adaptation to arise on rapid timescales independently in different hosts (Zhao et al. 2019; Torrillo and Lieberman 2024), without requiring recombination to spread adaptations so broadly across individuals. This scenario would result in ‘widespread parallelism’, in which no single haplotype reaches high frequency (Fig 1C). That we nonetheless observe gene-specific sweeps raises the question: do these sweeps bear signatures of slow or rapid adaptation? In the case where evolution is mutation-limited, an adaptation may take a long time to arise, and thus “hard sweeps” are expected. In this scenario, an adaptive mutation arises in a single host and transfers across individuals, resulting in a single adaptive haplotype rising to high frequency in the broader population (Fig 1A) (Pennings and Hermisson 2006). By contrast, when mutation is rapid, “soft sweeps” are expected. In this scenario, many adaptations may arise repeatedly in a short time span in distinct hosts that then spread, resulting in multiple adaptive haplotypes at high frequency in the broader population (Fig 1B) (Hermisson and Pennings 2005; Pennings and Hermisson 2006). Soft sweeps have already been invoked in human-associated and environmental microbial populations to explain the unexpected preservation of nucleotide diversity and the presence of multiple haplotypes or independent origins during adaptation (Derrick et al. 2024; Birzu et al. 2023; Bendall et al. 2016; Liu and Good 2024; Woods et al. 2020; Croucher et al. 2014). However, whether soft sweeps represent a common or even dominant mode of adaptation in the human gut, driving the spread of broadly beneficial alleles across populations, remains to be seen. Given the rapid influx of new mutations arising on distinct strain backgrounds across individuals’ microbiomes, we hypothesize that adaptation across human gut microbiomes will proceed predominantly through soft rather than hard sweeps driven by rare singular origins.

**Fig 1.**
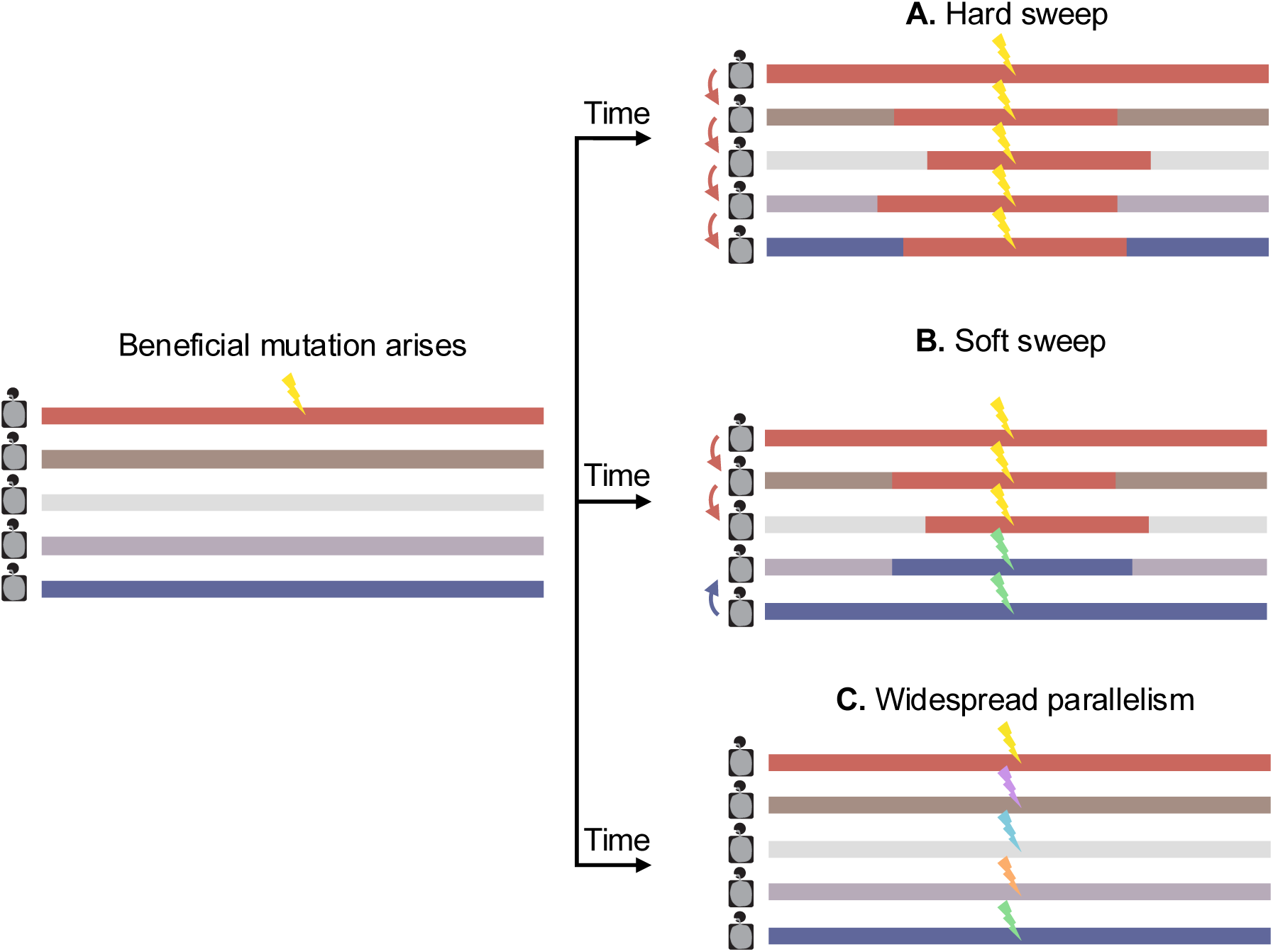
Signatures of adaptation in commensal gut species. After a beneficial mutation arises in a strain of one host’s microbiome, the spread of the adaptive mutation across multiple strains via recombination may result in a gene-specific selective sweep. Horizontal lines represent distinct genomes. Shared haplotypes have the same color. Thunderbolts represent adaptations. Adaptations in different colors have distinct origins. (**A**) The spread of a single beneficial origin across hosts results in a hard sweep, where a single haplotype is expected to reach high frequency. (**B**) When mutation is rapid, multiple origins of the beneficial mutation may sweep to high frequency on distinct haplotype backgrounds, resulting in a soft sweep. In this scenario, there are two origins sweeping. (**C**) However, if mutation is very rapid, then there may be a high number of unique origins of the adaptive mutation, which we term “widespread parallelism.” In this case, no haplotype is expected to reach high enough frequency for the adaptation to be detected as a sweep.

Here we analyze a panel of 693 hosts spanning North America, Europe and China to assess the prevalence of selective sweeps spreading across gut commensals and ascertain whether sweeps are hard or soft. First, confirming previous results (Wolff and Garud 2026), we find selective sweeps are pervasive across 10 common gut commensals analyzed. Second, contrary to the expectation of ubiquitous soft sweeps, we find that hard sweeps are also common. To reconcile this unexpected result, we consider the adaptive mutation rate required to observe sweeps in a metapopulation of gut microbiomes. Even with rapid recombination rates, we find that adaptation at single sites should result only in widespread parallelism, and consequently, that hard or soft sweeps must arise from complex, difficult-to-mutate adaptative architectures such as epistasis or adaptive introgression. Our results suggest that, contrasting with a picture of rapid adaptation at single sites, broadly shared recombination-mediated adaptations are often deeply mutation-limited, and that timescales of evolutionary innovation in the human gut may be far longer than previously appreciated.

## Results

### Detection of recombination-mediated selective sweeps in bacteria

To detect gene-specific selective sweeps spreading via recombination, we use a haplotype homozygosity statistic, H12, defined as follows:

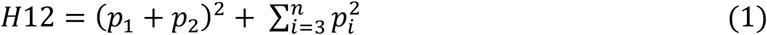

where *p*_i_ represents the frequency of the i^th^ most common haplotype among *n* haplotypes in a genomic window (Garud et al. 2015). By combining the frequencies of the two most common haplotypes in the first term, either one or multiple haplotypes at high frequency should similarly elevate H12, making the statistic capable of detecting both hard and soft sweeps. While H12 detects sweeps regardless of type, distinguishing hard from soft sweeps requires a complementary analysis, which we apply later.

While H12’s ability to detect hard and soft sweeps has been demonstrated in multiple eukaryotes, including *D. melanogaster* (Garud et al. 2015), domestic dogs (Schlamp et al. 2016), and many other species (Bolognini et al. 2024; Fuller et al. 2020; Frantz et al. 2015), it remains untested whether H12 can detect known selective sweeps in recombining bacterial populations. To evaluate H12 in this context, we applied H12 to the recombining bacterial pathogen, *Clostridiodes difficile*, with known targets of adaptation at several virulence-related loci, including recombination-mediated diversifying selection at the *tcdB* gene encoding the Toxin B virulence factor (Shen et al. 2020; Mansfield et al. 2020) and the S-layer cassette gene cluster (Dingle et al. 2013; Lynch et al. 2017; Yahara et al. 2016). Additionally, *C. difficile* harbors putative adaptations at the F1 and F3 flagellar operons, whose genes are important for gut adherence and pathogenicity (Steinberg and Snitkin 2020; Tasteyre et al. 2001).

Since H12 is applied in sliding windows along the genome, we identified species-specific window sizes in which local signals of excess homozygosity can be distinguished from the genomic baseline. Specifically, we identified window sizes that correspond on average to a length scale on which genome-wide linkage disequilibrium (LD) decays (and H12 should be low), but are not so large that elevated haplotype homozygosity arising from selective sweeps cannot be detected (Fig 2A, S5 Text). Additionally, we control for population structure, which could otherwise result in confounding signatures of elevated haplotype homozygosity (S3 Text).

**Fig 2.**
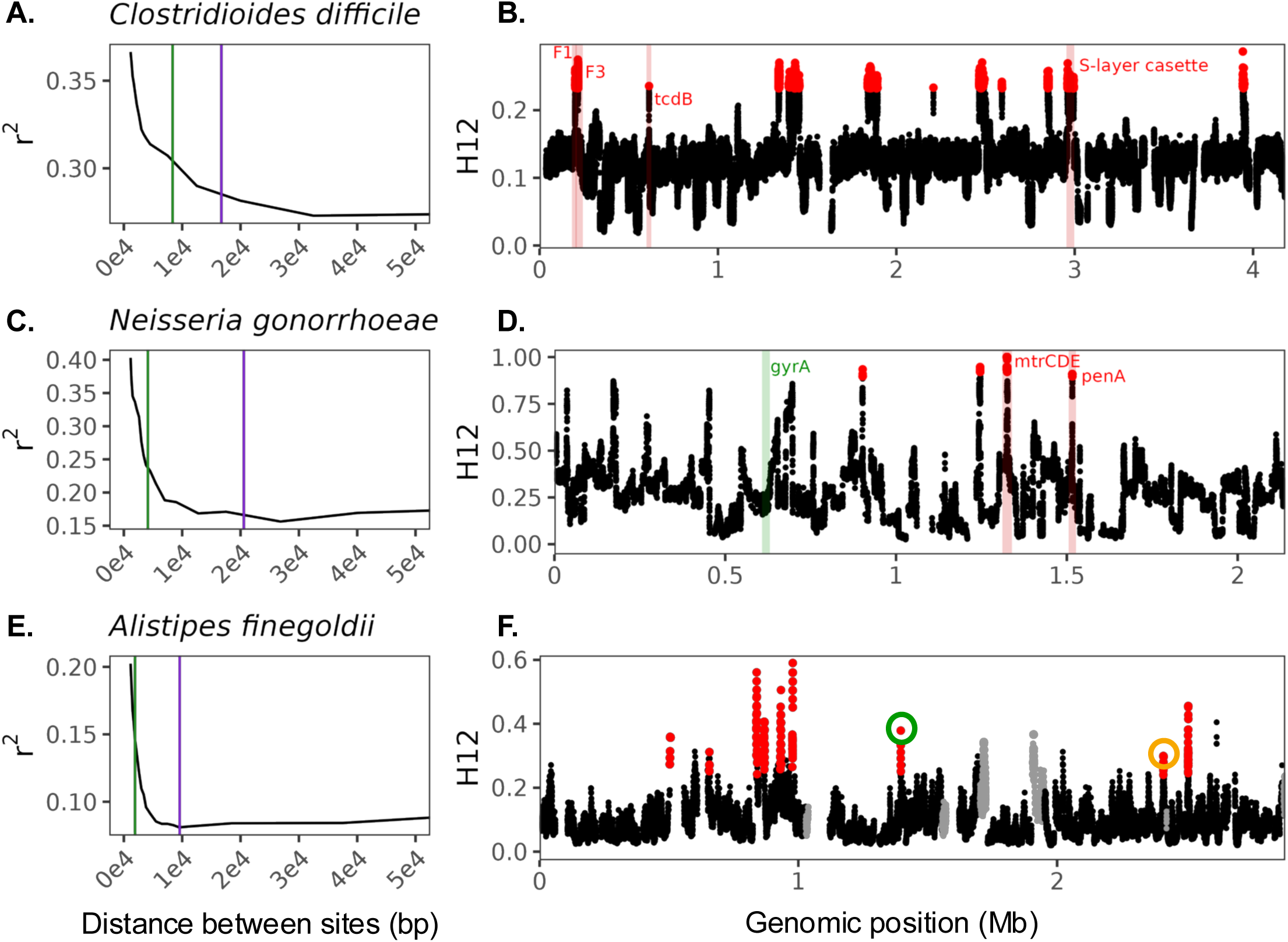
Scans for elevated haplotype homozygosity reveal selective sweeps in bacteria. Decay of linkage disequilibrium (r^2^) in (**A**) *C. difficile* (n=243 isolates), (**C**) *N. gonorrhoeae* (n=186), and (**E**) *A. finegoldii* (n=64) after population structure control (S3 Text). The green vertical line denotes the inferred mean tract length, and the purple vertical line, a multiple of the inferred tract length, denotes the window length used to run the H12 scan. An H12 scan of (**B**) *C. difficile* and (**D**) *N. gonorrhoeae*. Outlier windows in the scan with H12 values exceeding the 99th percentile are shown in red. (**F**) An H12 scan of *A. finegoldii*. Putative sweeps with H12 values inconsistent with neutrality are highlighted in red. H12 windows in regions with low recombination rates are grey dots (S5 Text). Orange and green circles indicate sweeps highlighted in Fig. 3 for hard vs soft inference.

We applied H12 to 243 isolates of *C. difficile* obtained from the Unified Human Gastrointestinal Genome (UHGG) collection (Almeida et al. 2020) (S1 Text), revealing sharp elevations in haplotype homozygosity at multiple loci when compared to the genomic baseline (Fig 2B). As nearby H12 windows may belong to the same sweep, we grouped adjacent windows with elevated H12 values into a ‘peak’ (S5 Text). Two prominent peaks coincide with *tcdB* and the S-layer cassette (Fig 2B), consistent with recent selective sweeps at these loci. Distinct peaks in H12 are also evident at the F1 and F3 flagellar operons, suggesting independent sweeps in each operon, a finding supported by a previous study showing unique haplotype structures in F1 and F3 (Steinberg and Snitkin 2020). Given our ability to recover known targets of positive selection at virulence loci, we conclude that H12 is sensitive to detecting known selective sweeps in *C. difficile*.

To further test the ability of H12 to detect known recombination-mediated selective sweeps in bacteria, we next applied H12 to 186 *Neisseria gonnorhoeae* isolates from the Gonococcal Isolate Surveillance Project (GISP) with genomic assemblies from Grad et al. 2014 (Haskin et al. 2025) (S1 Text). *N. gonorrhoeae* has positive controls of recombination-mediated sweeps at the well-characterized antimicrobial resistance loci *mtrCDE* (encoding mtrCDE efflux pump) and *penA* (encoding penicillin binding protein 2) (Grad et al. 2014; Wadsworth et al. 2018). Additionally, *gyrA* is a well-known target of adaptation to antimicrobial resistance to fluoroquinolones, but may arise predominantly through mutation as opposed to recombination, as strong resistance is conferred through single mutations at any of multiple sites (Belland, Morrison, and Huang 1994; Grad et al. 2016). Application of H12 revealed peaks at *penA* and the *mtr* operon (Fig 2C, 2D). However, we do not observe elevated haplotype homozygosity at *gyrA*, which we later show is consistent with a model of widespread parallelism at this locus (Fig 1C), in contrast with the other loci highlighted above.

### H12 scans in gut commensals reveal abundant selective sweeps

Having confirmed H12’s ability to detect adaptation in well-characterized pathogenic bacteria with known targets of positive selection, we next applied H12 to human gut commensal species from a set of metagenomic samples from 693 individuals from North America, Europe and China (Lloyd-Price et al. 2017; Korpela et al. 2018; Qin et al. 2012; Xie et al. 2016) that we previously collated and analyzed (Garud et al. 2019). As H12 is a haplotype-based statistic, we inferred strain haplotypes from individual metagenomes (S2 Text). We first aligned shotgun reads to a reference database to extract single nucleotide variants for each species in each host (Nayfach et al. 2016). When a microbiome harbors a single dominant strain of a species, that strain’s haplotype can be ‘quasi-phased’ such that major alleles within a host’s microbiome can confidently be assigned to the strain (Garud et al. 2019). Using this approach, we obtained 3316 quasi-phased haplotypes across many prevalent bacterial species in 693 individuals.

We applied H12 to the 28 species with at least 40 quasi-phased haplotypes. To assess peak significance, we simulate neutral distributions of H12 matched to the empirical baseline distribution of each species. As this distribution may differ from species-to-species, we conduct separate simulations for each species, matched to its estimated nucleotide diversity, recombination rate, and tract length (S4 Text). For all subsequent analyses, we focused on the 10 species for which we could assess peak significance against simulated neutral H12 distributions matched to the data (S5 Text, Fig S6). We found a total of 55 selective sweeps (Fig 2F, Fig S6), with a mean of 5.5 sweeps per species, in line with recent results discovering gene specific sweeps from gut bacteria (Wolff and Garud 2026). Together, these results are indicative of pervasive selective sweeps spreading across hosts in multiple gut commensal species.

### Prevalence of hard and soft sweeps in gut commensals

To gain an intuition whether the identified sweeps are hard vs. soft, we visually inspected haplotypes in windows with the highest H12 values at each identified sweep. This revealed in some instances one haplotype at high frequency, consistent with hard sweeps (Fig 3A), and in other instances multiple haplotypes at high frequency, consistent with soft sweeps (Fig 3C). The elevation in haplotype frequency in these windows is not observed in random windows in the genome or in simulations of neutrality. Instead, these windows visually resemble simulated hard and soft selective sweeps (Fig 3B, 3D, Fig S7).

**Fig 3.**
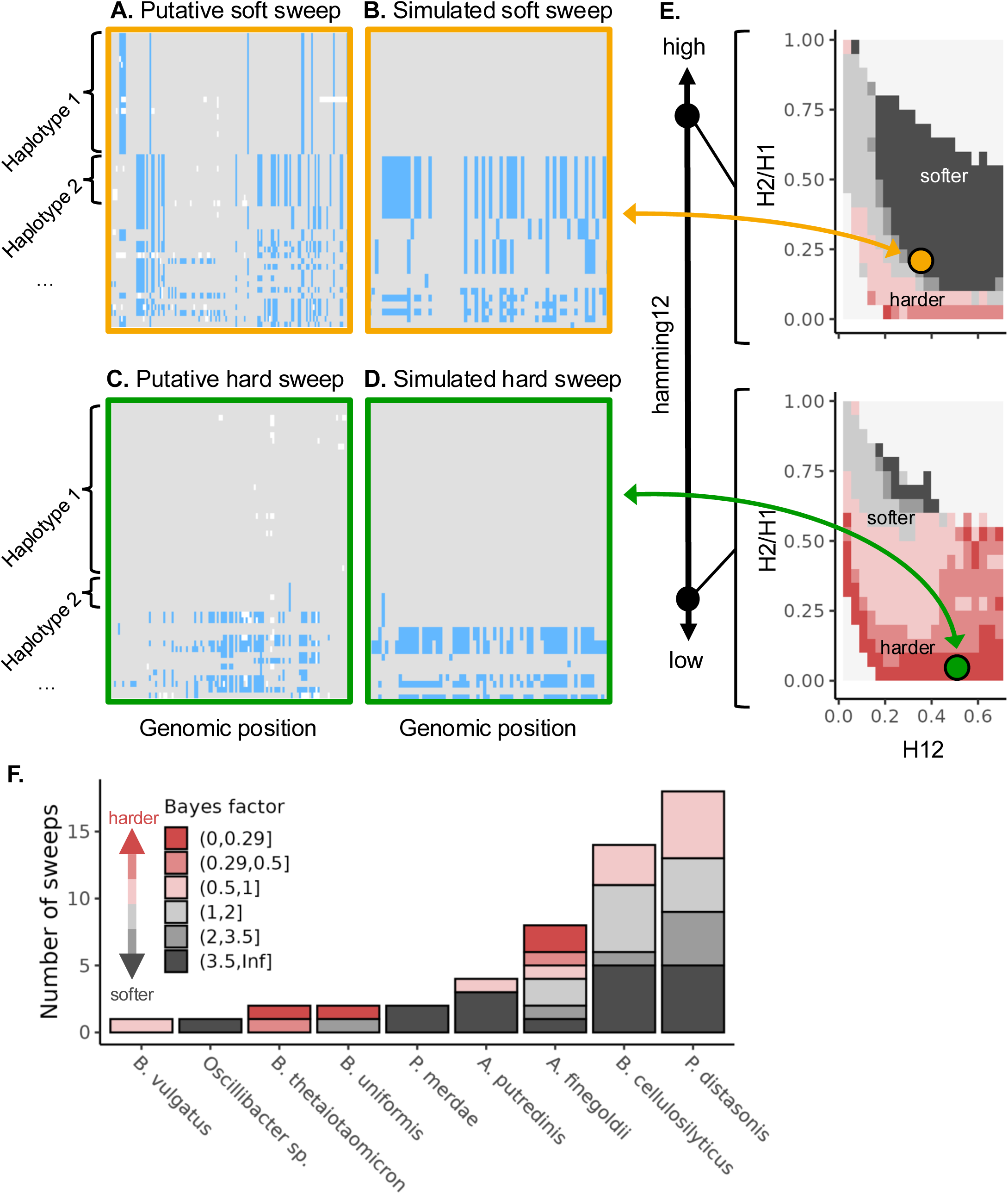
Distribution of hard and soft sweeps in gut commensal species. Selective sweeps are classified as hard or soft using an approximate Bayesian computation approach. Haplotype plots of (**A**) a putative soft sweep and (**C**) a putative hard sweep from the H12 scan in Fig 2 are compared to (**B**) a simulated soft sweep and (**D**) a simulated hard sweep whose values of H12, H2/H1 and hamming12 match the observed values at the putative sweep. (**E**) Bayes factors comparing support for a soft vs. hard sweep model as a function of H12, H2/H1 and hamming12 matching the putative sweep being analyzed. Irrespective of the hamming12 value, larger H2/H1 values better support soft sweeps. The putative soft sweep has a high Bayes factor of 10.25, with strong evidence for a soft sweep (orange dot), while the putative hard sweep has a low Bayes factor (0.17), with strong evidence for a hard sweep (green dot). (**F**) Distribution of Bayes factors among all inferred sweeps across 10 species. BFs <1 indicate support for hard sweeps and BFs =1 indicate support for soft sweeps.

To confirm this intuition, we next statistically inferred whether haplotype signatures at the selective sweeps identified are more likely to arise from hard vs soft sweeps. To do so, we used approximate Bayesian computation (ABC) to compute the likelihood that a sweep is hard or soft by comparing three summary statistics of haplotype structure – H12, H2/H1, and hamming12 – measured from simulated hard and soft sweeps to those measured in the data (S6 Text). In brief, while H12 captures the overall magnitude of haplotype homozygosity, H2/H1 captures the proportion of homozygosity without the most common haplotype and is expected be elevated under soft sweeps relative to hard sweeps (Garud et al. 2015). Introduced in this paper, hamming12 measures the sequence divergence between the two most frequent haplotypes. Hamming12 is expected to be large for soft sweeps because high-frequency haplotypes originating from independent strain backgrounds will be divergent. While hamming 12 should also be large for a hard sweep when the adaptive haplotype is diverged from any non-adaptive haplotype, hard sweeps whose haplotypes have broken due to occasional passenger mutations or recombination events may exhibit low hamming 12 values, allowing us to rule out any hard sweep that may resemble a soft sweep in terms of haplotype frequency. We show that the inclusion of hamming12 with the two other summary statistics improves the ability to discriminate between hard and soft sweeps (Fig S10).

To perform ABC, we extend the species-specific simulations described previously (and see S4 Text) to simulate selective sweeps. We draw the selection coefficient (*s*), partial frequency of the sweep (*PF*), and number of generations since selection ceased (*T*) from uniform prior distributions as follows: *s* ∼ Unif(0,0.2), *PF* ∼ Unif(0,1), and *T ∼* Unif{0,200}. The number of origins in a selective sweep is determined by the population-scaled mutation rate at the adaptive locus, Θ_A_. A value of 1 is expected to introduce on average 1 mutation per generation in a population, and will almost exclusively generate soft sweeps (Hermisson and Pennings 2005; Pennings and Hermisson 2006). We simulated 300,000 soft sweeps with Θ_A_ of 1 for each species. An equal number of hard sweeps are simulated by seeding only one beneficial mutation at the adaptive locus. We then compared an empirical sweep’s values of H12, H2/H1 and hamming12 to all simulated values, retaining simulations within a small normalized Euclidean distance to the empirical values (S6 Text). We then calculated the Bayes factor (BF), representing the relative strength of evidence for one model over the other, as the ratio of the number of matching simulated soft vs hard sweeps.

Fig 3E shows the range of BFs for a grid of H12 and H2/H1 values, conditional on hamming12 values for two putative sweeps identified in *A. finegoldii* in Fig 2 (the full distribution is shown in Fig S9). Of the two sweeps highlighted in Fig 2F, one has substantial support for being hard with a BF=0.17, while the other has strong support for being soft with a BF=10.25 (Fig 3E). More broadly, we compute BFs for all sweeps inferred across all 10 species analyzed that are supported by at least 30 simulations (Fig 3F). There is a wide distribution of BFs among all species, spanning very strong support for hard sweeps to very strong support for soft sweeps. In total, 17 sweeps have support for hard sweeps (BF <1) and 36 sweeps have support for soft sweeps (BF>1) (Fig 3F). Of these, 4 sweeps have substantial evidence for hard sweeps (Bayes factor < 0.29) and 18 sweeps have substantial evidence for soft sweeps (Bayes factor > 3.5).

The inference of several hard sweeps in our data is at odds with expectations for ubiquitous soft sweeps when mutational inputs are estimated to be on the order of a billion per day per human microbiome (Zhao et al. 2019). To better understand this result, we next consider what population genetic mechanism would permit a hard sweep to spread across multiple human gut microbiomes given such rapid mutational input.

### Sweeps spreading across human gut microbiomes cannot be driven by *de novo* mutations at a single site

Hard sweeps, or any gene-specific sweep for that matter, can only arise via recombination between gut commensal strains and eventual migration of those strains into recipients’ gut microbiomes. If *de novo* mutation is sufficiently rapid, many gut microbiomes will evolve adaptations independently before acquiring them via migration and recombination, resulting in widespread parallelism (Fig 1C). Thus, we expect to observe hard sweeps only if migration and recombination rates sufficiently outpace mutation rates.

To assess the range of recombination and mutation rates that would allow for sweeps to spread across human gut microbiomes, we considered a stochastic model of the spread of adaptations across a population of hosts, each harboring a single strain. In this model, selection acts on one strain at a time, at which point the strain may either *de novo* mutate an origin of the adaptive mutation or obtain the adaptation from another host’s strain via migration and subsequent recombination.

The rate at which a strain obtains an adaptive allele via migration and recombination vs *de novo* mutation is:

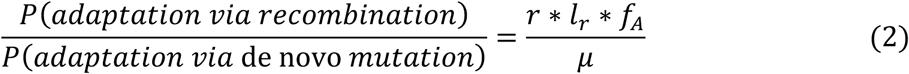

where *r* is the per-base pair recombination rate, *l_r_* is the average homologous tract length exchanged, and *μ* is the per-base pair mutation rate. Migration is included implicitly in the recombination rate parameter because the empirical recombination rates considered below were estimated across host microbiomes. The quantity *f*_A_ describes the fraction of strains carrying the adaptation. Thus, the first strain to adapt (when *f*_A_ = 0) has zero odds of acquiring the adaptation via recombination, and instead mutates an adaptive origin with 100% probability. Every subsequent strain has increasing odds of adapting via recombination because *f*_A_ increases over the course of the sweep (Fig S11). To assess a range of mutation and recombination rates, we vary r ∗ *l_r_* / µ, where large values (>>1) will result in fewer origins at higher frequencies (e.g. harder sweeps) due to recombination outpacing mutation, and very small values <1 will result in many origins at lower frequencies (e.g. soft sweeps) due to mutation outpacing recombination.

With the above in mind, the stochastic model describes a Chinese restaurant process (CRP) with parameter (N ⋅ µ) / (r ⋅ *l_r_*) (S7 Text). In this model, N is the census population size at the time of the sweep. We used analytical expressions from the CRP to calculate the expected number of origins and frequencies of origins across a large range of *r* ∗ *l_r_* / *μ*, with N fixed at 10^8^, though we note our results do not qualitatively change when N changes by orders of magnitude (Fig S12).

Empirical estimates of *r* ∗ *l_r_* / *μ* vary from 10^2^ – 10^3^ (Liu and Good 2024), which means that recombination should outpace mutation. However, despite recombination seeding the adaptation in the vast majority of hosts in this parameter range, our model predicts sweeps having 10^5^ – 10^6^ origins (Fig 4A, Fig S11). With so many origins, even the most abundant origins cannot surpass a frequency of 10^-4^ (Fig 4B). In this scenario, the probability of any two hosts sharing an origin is < 2.47e-5, implying that in a random sample of a population of strains (across hosts), virtually all will harbor a unique origin. Any haplotype sharing between strains will be incidental.

**Fig 4.**
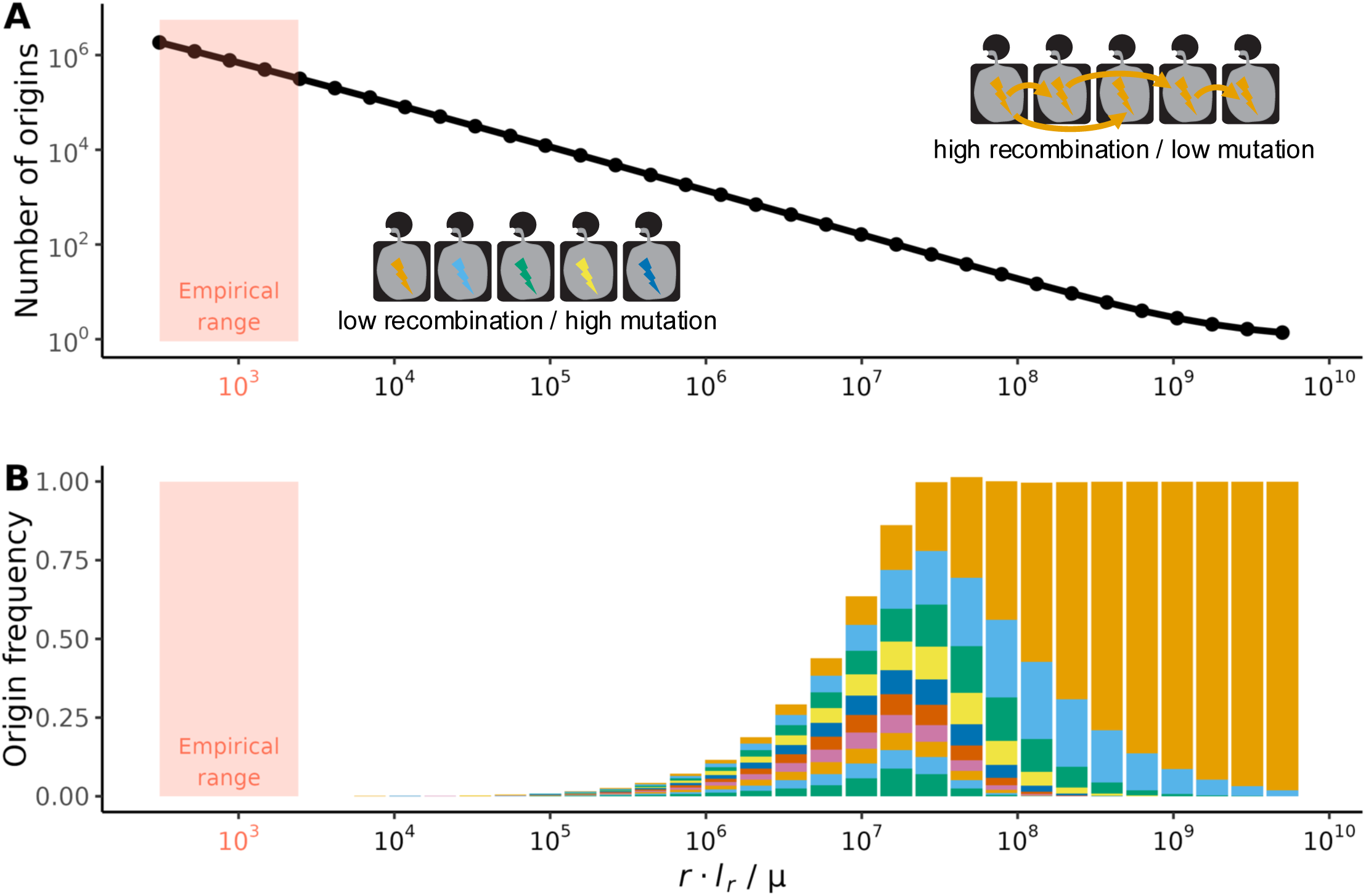
Expected number of origins and frequencies of the 10 most common origins as a function of the ratio of recombination vs mutation rate. We modeled the spread of an adaptation across hosts with a stochastic model. In this model, strains can either mutate an adaptation *de novo* with rate μ or acquire an adaptation from another strain via recombination with rate r * l_r_. The pink vertical band denotes the range of recombination-to-mutation rate ratios estimated in common commensal gut bacteria in (Liu and Good 2024). (**A**) Expected number of origins of the adaptation as a function of r * l_r_ / μ in a population of 10^8^ hosts. (**B**) Expected frequencies of the 10 most-frequent origins in the same model. The height of each color band denotes the frequency a different origin. For example, in the last band, a single origin is at almost 100% frequency in the population.

By increasing *r* ∗ *l_r_* / *μ* by 4 – 5 orders of magnitude, the number of origins decreases to 10^1^ – 10^2^. In this range, the frequencies of the 10 most common origins are collectively close to 1, with nearly every host bearing one of these 10 origins. In this scenario, the probability of any two hosts sharing an origin is as high as 0.087, implying that in a random sample, many hosts should share an origin of the adaptive allele. Finally, by 6 orders of magnitude, the expected number of origins is on the order of 1 and the expected frequency of the most frequent origin is near 100 percent. This implies that nearly every host bearing the adaptation should share the same origin.

It is important to note that the model we implement results from us making several conservative assumptions that increase the probability of a hard sweep (S7 Text). Despite this conservativeness, the model struggles to produce hard sweeps in the empirically-estimated range of recombination and mutation, suggesting that widespread parallelism should instead be predominant.

This makes our observations of hard sweeps all the more surprising. In order to observe hard or, for that matter soft sweeps, a 4-to-6 order of magnitude increase in the empirical recombination-to-mutation ratio is required. One way this increase can be achieved is if the recombination rate is far greater than measured from data. However, instead of a physiologically unrealistic increase in the estimated recombination rate, we hypothesize that the adaptive mutation rate at sweeps spreading across hosts is significantly lower than expected for single base pair sites.

Such a reduction in the adaptive mutation rate may be biologically plausible if adaptation proceeds through complex, multisite adaptive architectures. For example, structural or accessory elements in a bacterial genome likely cannot *de novo* mutate on the timescale of a sweep even in the largest of bacterial population sizes. Alternatively, an adaptation may require multiple epistatically interacting mutations to generate the adaptive benefit. We show in S7 Text that a simple model of pairwise reciprocal sign epistasis, where two mutations that are strongly deleterious alone but are beneficial together, is consistent with this hypothesis and may result in sweeps across hosts.

### Gene specific sweeps bear genomic signatures of complex adaptive architectures

Our model predicts that adaptations sweeping across gut microbiomes must harbor complex, multi-site adaptations. To determine if this prediction is borne out by the data, we investigated whether adaptations detected with H12 exhibit genomic signatures of complex multisite adaptive architectures. First, to gain intuition on these genomic signatures in known controls, we used *N. gonorrhoeae* as a case study, as the genetic architecture of its resistance to various antibiotics is well-characterized (Wadsworth et al. 2018; Belland, Morrison, Ison, et al. 1994; Tomberg et al. 2010).

In *N. gonorrhoeae*, complex multi-site adaptations at the *penA* gene and *mtr* operon have been shown to arise from the introgression of ‘mosaic’ alleles from other, distantly related commensal *Neisseria* species. As shown experimentally, combinations of mosaic alleles at these loci epistatically interact to confer resistance (Wadsworth et al. 2018; Tomberg et al. 2010). In contrast, *gyrA*, a well-known target of adaptation conferring antimicrobial resistance to fluoroquinolones, is not a multi-site adaptation; only a single mutation is required to confer resistance, which typically evolves rapidly through *de novo* mutation, and seldom through recombination (Belland, Morrison, and Huang 1994; Grad et al. 2016).

The H12 scan of *N. gonorrhoeae* (Fig 2) revealed high H12 values at the *penA* and *mtr* loci. This finding supports the prediction that haplotypes bearing complex adaptations will recombine onto multiple, distinct genomic backgrounds. By contrast, there is no elevation of H12 at the *gyrA* locus, despite the adaptive resistance allele (S to F at position 91) being present in 77% of the strains analyzed. This lack of signal is consistent with widespread parallelism (Fig 1C), in which no sweep is observed because haplotypes cannot reach high frequencies across hosts.

Additional genomic signals at *penA* and *mtr* provide further evidence that these adaptations are inconsistent with sweeps arising from adaptive *de novo* mutation at single sites. Nucleotide diversity (π) is sharply elevated at both loci, consistent with the introduction of novel variation from hybridizing species (Unitt et al. 2024; Wadsworth et al. 2018) (Fig S13). At *penA*, π reaches the highest quantile among all windows in the genome, at *mtr, π* reaches a high quantile of 0.95, while at *gyrA, π* reaches the 0.59 quantile, near the genome-wide median (Fig 5A). This signature of elevated π at *penA* and *mtr* is consistent with their introgressed origin and inconsistent with a depression in π expected for a classic selective sweep arising from a single point mutation (Nielsen 2005).

**Fig 5.**
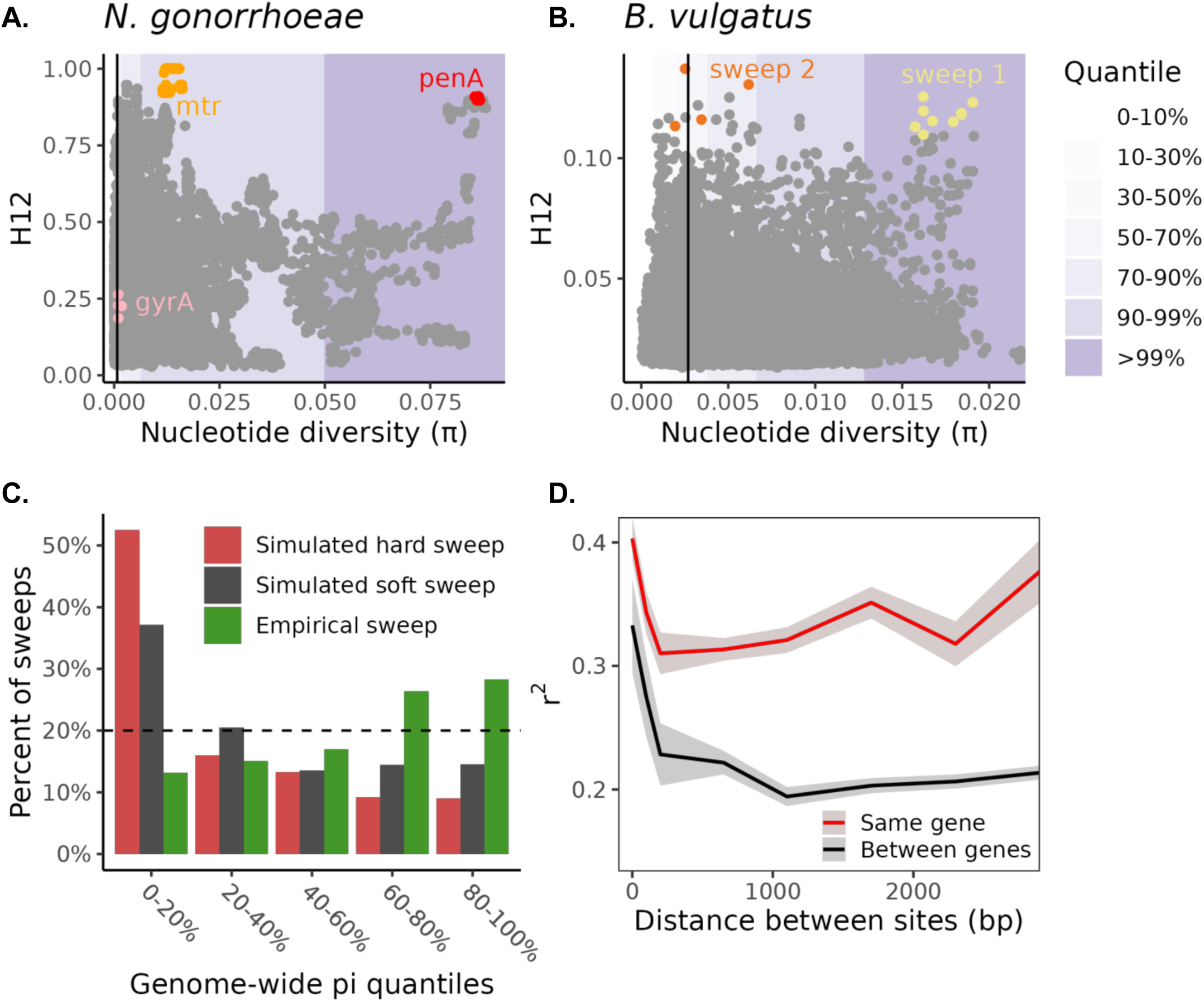
Evidence for complex multisite adaptations at selective sweeps. H12 vs. π in (**A**) *N. gonorrhoeae* and (**B**) *B. vulgatus.* Each point represents an analysis window in the H12 scan. Background shading indicates a window’s quantile of π relative to the genome-wide distribution of π. Black vertical lines denote the genomic median of π. Highlighted for *N. gonorrhoeae* are known targets of selection and for *B. vulgatus* two putative sweeps, with one sweep attaining quantiles of π greater than 0.99. (**C**) Quantiles of π for empirical vs simulated sweeps. In the case of empirical sweeps, quantiles were calculated relative to the genome-wide distribution. In the case of simulations, quantiles were computed relative to neutral simulations. As a visual aid, we plot the 20% threshold to emphasize which bins are especially enriched or depleted in sweeps as compared to the expected genome-wide (or neutral) distribution of π. (**D**) Decay of LD (*r^2^*) in four sweeps with excess of LD within vs between genes (For full distribution, see Fig S18).

Additionally, Wadsworth et al. 2018 report high linkage disequilibrium (LD) over the length of *mtrD* and its associated promoter region followed by quick decay to genomic baseline, which they experimentally confirm corresponds to a region of strong allelic epistasis (Fig S14). A region of elevated LD in *penA* similarly aligns with experimentally confirmed epistasis in a resistant mosaic background, with LD quickly decaying at the gene’s boundaries (Fig S14) (Tomberg et al. 2010). The requirement of multiple synergistically interacting mutations to confer the adaptive benefit in epistatic regions is likewise definitively inconsistent with adaptation via a single *de novo* mutation. On the other hand, no elevation in LD is detected in *gyrA,* consistent with its high adaptive mutation rate and lack of synergistic epistasis (Fig S14).

Having examined signatures of nucleotide diversity and LD in *N. gonorrhoeae* controls, we next examined whether similar signals exist in the 55 sweeps identified via H12 in commensal gut species. We first calculated the distribution of π quantiles at these sweep locations relative to their species-specific genomic backgrounds. As an example, one sweep in *B. vulgatus* reaches the highest quantiles of π (>99%) among all windows in the genome, resembling the *penA* locus (Fig 5B, Fig S13). Across the 55 sweeps, the distribution of π quantiles was surprisingly broad, with elevated quantiles of nucleotide diversity when compared to the genomic background. In fact, more than 50% of empirical sweeps attained π quantiles between 60% and 100%. By contrast, simulations of hard and soft sweeps arising from single-site mutations showed the expected decrease in nucleotide diversity (enrichment of low π quantiles) (Fig 5C). In these scenarios, less than 20% and 30% of hard and soft sweeps, respectively, achieved quantiles between 60% and 100%. While it may be difficult to determine if any individual sweep is introgressed, this enrichment of high quantiles of π in the empirical data is highly inconsistent with classic sweeps arising from a single base pair mutation.

We then calculated LD between variants at all sweeps, conditioning on whether variants reside in the same gene or different genes, as several studies have noted that elevated LD within genes compared with between genes at matched distances is consistent with synergistic epistasis (Stolyarova et al. 2022; Callahan et al. 2011; Sandler et al. 2021). While the overall distribution of differences in LD within vs between genes was centered around zero, 21 of 55 sweeps exhibited higher LD within genes than between them (Fig S15). Four sweeps in particular had a pronounced increase in LD within vs between genes (more than 2 standard deviations above the mean), elevating LD values at all measured distances (Fig 5D, Fig S15). We note that while not all sweeps have elevated LD within vs between genes, in several cases LD is elevated over the length of multiple gene boundaries before dropping precipitously (Fig S16). Possibly, epistatic interactions can be maintained over multiple genes or whole operons rather than just one gene, leaving our approach underpowered to detect the full extent of epistasis in selective sweeps.

As an example, we examined the sweep in *B. vulgatus* shown in Fig 5B exhibiting excess π. In this case, a block of LD is elevated within the boundaries of a predicted polysaccharide utilization locus composed of several genes (PUL) (Terrapon et al. 2018), beyond which it quickly decays to genomic baseline (Fig S13, Fig S16). As PUL genes are expected to be tightly co-regulated and co-expressed (Grondin et al. 2017), one plausible explanation is that epistatic loci spanning the length of the PUL have contributed to the complex multisite adaptive architecture of this sweep.

In summary, we found that multiple genomic signatures, including elevated H12, elevated nucleotide diversity, and increase in within-gene vs between-gene LD are inconsistent with sweeps arising from single site mutation. We conclude that mechanisms consistent with complex adaptation, such as introgression or epistasis involving multiple alleles, may be widespread among gut commensal species, even when processes such as simple pairwise reciprocal epistasis may be sufficient to enable gene-specific sweeps (S7 Text).

## DISCUSSION

In this work, we investigated whether gene-specific selective sweeps spreading across human gut microbiomes arise via hard vs. soft selective sweeps. We expected to see exclusively soft sweeps, in which multiple origins of the same adaptation sweep to high frequency, consistent with the rapid mutational input to the human gut microbiome (Zhao et al. 2019). Instead, we found a mixture of hard and soft sweeps. Our modeling reveals that sweeps of any kind only become probable when adaptations are difficult to mutate *de novo*. Thus, we propose that gene-specific sweeps spreading across hosts harbor complex adaptive architectures characterized by multisite allelic variation, such as epistasis and structural variation, and we find genomic signatures in the vicinity of sweep regions supporting this.

Examples of complex adaptive architectures are well established among pathogens but their prevalence and targets in gut commensals have not been characterized thus far. For example, in *Neisseria gonorrhoeae* the multiple transferable resistance efflux pump confers drug resistance via complex epistasis through domain mosaicism at the *mtr* operon (Wadsworth et al. 2018), and resistance at the penicillin binding protein 2 arises similarly at *penA* (Tomberg et al. 2010). The pathogen *Clostridiodes difficile* has evolved adaptations at a toxin B virulence factor, *tcdB*, also via domain mosaicism whereby recombination has brought together pre-adapted domains with varying levels of pathogenicity (Janezic et al. 2020). Epistatic interactions among four genes enabled a strain of *Pseudomonas aeruginosa* to become a dominant pathogen in the human lung (Damkiær et al. 2013). A structural insertion sequence, ISL3, in *Enterococcus faecium* may provide adaptive advantage in hospital-associated infections (Grieshop et al. 2026). In influenza A, surface proteins have evolved epistatic adaptations to escape host immunity and drugs (Kryazhimskiy et al. 2011). These and other examples illustrate that complex adaptations abound in microbial pathogens (Cummins et al. 2026; Wong 2017). Our work reveals that complex adaptations may not be specific to pathogens and are a common feature of commensal gut microbiota, presumably in response to common selection pressures of the gut, such as diet or the host immune system.

Our stochastic model showed that single site adaptations are consistent only with widespread parallelism, in which independent adaptive origins are so abundant that none reach high frequency (Fig 1C). Assuming single site adaptations abound and complex multisite adaptations are relatively rarer, widespread parallelism should be the norm. Contrary to this expectation, gene-specific sweeps seem to be common (Fig 3, Wolff and Garud 2026), and across-host parallelism has been characterized as modest thus far (Lieberman 2022; Scanlan 2019; Zhao et al. 2019; Garud et al. 2019; Chen and Garud 2022). Therefore, an important direction for future research is to quantify the relative extent by which bacteria respond to selection via complex, multisite variation versus single-site mutations. Such information would have broad implications for the dynamics and spread of clinically relevant adaptations. For example, antibiotic resistance mutations that can rise to high frequency within hosts easily in a matter of days might render the drug ineffective for pathogens but also allow beneficial gut commensal microbes to survive. Meanwhile, difficult-to-evolve antibiotic resistance adaptations may retain the drug’s effectiveness in eliminating pathogens, but result in more extensive collateral damage to beneficial microbiota.

Our previous work leveraged signatures of deleterious mutations hitchhiking with the beneficial allele to find gene-specific selective sweeps in the microbiome (Wolff and Garud 2026). Now, our results suggest that gene-specific sweeps harbor complex adaptations, which was not explicitly accounted for in Wolff. However, we expect the two models are completely compatible, as deleterious mutations are likely hitchhiking alongside beneficial alleles regardless of whether those alleles involve simple or complex architectures. Gene-specific sweeps in the microbiome therefore likely carry a mixture of complex adaptive variation and deleterious alleles hitchhiking with the selected haplotype.

Understanding the complex adaptive architecture underlying gene-specific selective sweeps is an exciting avenue for future research. Several key questions remain open. First, what role does epistasis play in driving adaptation? This topic is just beginning to be systematically explored (Arnold et al. 2018), and we predict that pairwise interactions between as few as two loci may be sufficient to produce sweeps (S7 Text). Yet our data suggest potentially much more complex epistasis, as we observe elevated linkage disequilibrium across multiple genes and operons, as seen in a polysaccharide utilization locus in *B. vulgatus* (Fig S13, Fig S16). Second, could structural variants be a key driver of adaptation? Mobile genetic elements (Fogarty et al. 2024; García-Bayona et al. 2024; Sheahan et al. 2024; Hehemann et al. 2010), insertion/deletions (potentially of whole pathways) (Zeevi et al. 2019; Sheahan et al. 2024), and chromosomal inversions (Jiang et al. 2019; Chanin et al. 2024) could all be candidates. Long read sequencing data may be particularly valuable for detecting such elements. Finally, the sources of adaptive variation remain unresolved. Do beneficial mutations typically arise within a population and then sweep locally, or are they commonly introgressed from more distantly related lineages? We observe signals of elevated nucleotide diversity in our sweeps (Fig 5), suggesting that adaptive introgression is widespread. New statistics will be crucial for disentangling these and other sources of adaptive variation.

If all adaptations at gene-specific sweeps are mutationally constrained, then the mixture of hard and soft sweeps we observe raises the question of what conditions might favor one vs the other. One possibility is there is substantial heterogeneity in the adaptive mutation rate at different adaptive loci, which would result in more accessible adaptations giving rise to soft sweeps and less accessible mutations to hard sweeps. Another possibility is that limits to migration determine the softness of sweeps, with higher rates of migration giving rise to hard sweeps, and lower rates resulting in soft sweeps (Ralph and Coop 2010; Paulose et al. 2019). Heterogeneity in migration rates may arise from biological properties of the bacterial species (e.g. tolerance to oxygen) (Browne et al. 2016) or properties of the host population (e.g. highly isolated group) (Carter et al. 2025), and their interactions with how selection pressures are distributed. Disentangling these effects might prove useful in predicting which species are especially predisposed to adapt via complex architectures, and which might exhibit local signatures of adaptation.

In sum, our study reveals that gene-specific selective sweeps spreading across human gut microbiomes comprise a mixture of hard and soft events, and that complex adaptive architectures are an essential feature of these sweeps. Our work highlights the importance of recombination-driven adaptation that may be difficult to generate by mutation alone. Concretely, the potential for an individual’s microbiome to rapidly respond to a selection pressure may depend on the reservoir of available adaptive material that the community at large provides. This insight may have broad implications for the development of fecal microbiome therapies, where donors harboring key beneficial variants could be valuable candidates for therapeutic transfer. More generally, continued efforts to characterize the tempo and mode of adaptation in the human gut microbiome promise to illuminate fundamental properties of microbial evolution in these complex communities.

## Acknowledgements

We thank Richard Wolff, Pleuni Pennings, Alison Feder for feedback on the manuscript, as well as all members of the Garud lab for helpful discussions and feedback on the manuscript. This work was funded by US National Institutes of Health National Institute of General Medical Sciences award R35GM151023, a National Science Foundation CAREER award (no. 2240098), and a Paul Allen Research Foundation grant to N.R.G., as well as National Institutes of Health National Human Genetics Research Institute award T32HG002536 supporting P.J.L.

## Data availability

Accession codes for the metagenomic data analyzed in this paper are as follows: PRJNA48479, PRJNA275349, PRJEB9576, PRJNA422434 and PRJEB24041. A full set of individual samples is included in Supplementary Table 1. For analyses involving *C. difficile* UHGG data, we downloaded alignments of MAGs and isolates, as well as accompanying genomic data files used to annotate sites and genes, from MGnify (https://ftp.ebi.ac.uk/pub/databases/metagenomics/mgnify_genomes/human-gut/) (Almeida et al. 2020). For analyses involving *N. gonorrhoeae*, we downloaded assemblies using accession number PRJEB2999 (Grad et al. 2014).

## Methods

All methodology can be found in the Supplemental Information File.

