## Supplemental Information for "Complex adaptive architectures constrain the pace of adaptations sweeping across human gut microbiomes"

#### Supplemental Methods

1. *Clostridioides difficile* and *Neisseria gonorrhoeae* data ................................................4

2. Metagenomic pipeline.....................................................................................5
   a) Overview
   b) Data
   c) Metagenomic processing with MIDAS
   d) Quasi-phasing
   e) Determining core genes

   a) Determining the window size for H12 scans
      Inferring the mean tract length
   b) Control for missing data
   c) H12 computed on synonymous sites carries greater power to detect sweeps
   d) Removal of regions with low recombination rate
   e) Calling significant H12 peaks

6. ABC classification of soft and hard sweeps .................................................................17
   a) Overview
   b) Summary statistics of haplotype structure
      H2/H1
      Hamming12

7. Stochastic model of a sweep across host microbiomes ..................................................20
   a) Overview

b) Model assumptions
c) Modeling the spread of an adaptation with the Chinese restaurant process
d) The expected number of origins and origin frequencies under the Chinese restaurant process
e) Approximating the waiting time for adaptations to arise under reciprocal pairwise epistasis

8. Genomic signatures of complex adaptations ............................................................... 27
a) Calculating nucleotide diversity quantiles

#### Supplemental Figures

Fig S1. Overview of H12 with downsampling ........................................................................28

Fig S2. Elevation of empirical distributions of H12 compared to neutral simulated distributions of H12 in *Alistipes finegoldii* ...............................................................................................29

Fig S3. Population structure correction improve the decay of linkage disequilibrium in *B. vulgatus* .........................................................................................................................30

Fig S4. Comparison of mean tract lengths $l_r$ and $l_r^{LD}$ among gut commensal species................31

Fig S5. H12 computed on synonymous variants has increased power when compared to H12 using synonymous and nonsynonymous variants....................................................................32

Fig S6. H12 scans in gut commensal species ..............................................................................33

Fig S7. Haplotype diversity in *A. finegoldii* at H12 peaks compared to simulated hard and soft selective sweeps...........................................................................................................35

Fig S8. Soft sweeps arising from SGV are still distinguishable from hard sweeps using hamming12 ...........................................................................................................................36

Fig S9. Distribution of Bayes factors across hamming12, H12 and H2/H1 for *Alistipes finegoldii* ...........................................................................................................................................37

Fig S10. hamming12 improves ABC classification at sweep posteriors....................................38

Fig S11. Adaptations are primarily obtained via recombination in a stochastic model of an across-host sweep.........................................................................................................................39

Fig S12. Number of origins and frequencies of sweeping haplotypes as a function of number of hosts in the population ....................40

Fig S13. Local patterns of H12 and nucleotide diversity at three regions in *N. gonorrhoeae* and one sweep in *B. vulgatus* ....................41

Fig S14. Local patterns of linkage disequilibrium at *mtr* and *penA* in *N. gonorrhoeae* ....................42

Fig S15. The difference in linkage disequilibrium AUC within genes vs. between genes for all gut commensal sweeps ....................43

Fig S16. The maintenance of LD over multiple gene boundaries in four gut commensal sweeps ....................44

**Supplemental Table Descriptions** ....................45

Table S1. H12 parameters for selection scans

Table S2. Simulation parameters for individual species

Table S3. H12 and summary statistics in sweep regions

Table S4. Summary statistics of haplotype structure and Bayes factors for sweeps

#### 1. *Clostridioides difficile* and *Neisseria gonorrhoeae* data

We downloaded SNP data from *C. difficile* isolates from UHGG v 1.0 (Almeida et al. 2020) (direct download link: https://ftp.ebi.ac.uk/pub/databases/metagenomics/mgnify_genomes/human-gut/v1.0/snv_catalogue/MGYG-HGUT-02369.tsv.tar.lz4). To annotate coding regions as synonymous and non-synonymous, we used custom scripts that used the reading frame coordinates to check for codon degeneracy. To correct for population structure, we used the approach described in S3 Text: "Population Structure Correction." Population structure correction resulted in 243 haplotypes of diverged and recombined strains. To choose an appropriate window size to apply H12, we followed the methodology described in S5 Text: "Determining the window size for H12 scans." All other H12 parameters, including the downsample fraction and missing data threshold, match the default for other gut commensal species (S5 Text: "Control for missing data" and Table S1). Virulence factor annotations were obtained by performing a BLAST search of the reference assembly against the VFDB database for *C. difficile* (Zhou et al. 2024; Altschul et al. 1990). We annotated *C. difficile* genes as virulence factors if more than 90% of codons matched the VFDB entry as per Wolff and Garud 2026.

For *N. gonorrhoeae,* we downloaded all high-quality isolates assembled in (Grad et al. 2014) (accession number PRJEB2999, and originally sourced from the Gonococcal Isolate Surveillance Project (GISP) (Haskin et al. 2025). We aligned all assemblies to the reference isolate FA0190 (RefSeq Assembly GCF_000006845.1) and called SNPs using the nucmer tool from MUMmer4 v 4.0.0rc1 (Marçais et al. 2018), following a similar approach as the UHGG v1 SNP calling pipeline (Almeida et al. 2020). We annotated SNPs as synonymous and nonsynonymous, corrected for population structure, and chose window sizes using the same procedures as described above, resulting in 186 strain haplotypes. The full H12 parameter set can be seen in Table S1.

#### 2. Metagenomic pipeline

##### a) Overview

All metagenomic data were processed using procedures described previously (Garud et al. 2019). In brief, since H12 requires phased haplotypes, we first map reads to reference genomes of commensal gut bacteria and then identify samples where a haplotype for a given species can be inferred with high accuracy. Below we describe the steps taken to process the data.

##### b) Data

We analyze a collection of metagenomic data collated and described in a previous study (Garud et al. 2019). This collection includes 693 subjects from the Human Microbiome Project (250), Xie et al (250), Qin et al (185), and Korpela et al (8) (Lloyd-Price et al. 2017; Xie et al. 2016; Qin et al. 2012; Korpela et al. 2018). As we do not investigate the longitudinal aspects of the data, we only use one metagenomic sample per subject (randomly choosing one sample if multiple have phaseable haplotypes).

##### c) Metagenomic processing with MIDAS

Using MIDAS v. 1.2 (Nayfach et al. 2016), we mapped metagenomic reads to a collection of universal single-copy genes to determine species presence and abundance, and subsequently inferred gene and SNV content for individual species by mapping to a collection of commensal reference determined to be present in the sample. Mapping parameters are as described in Garud et al. 2019. Species were deemed present by having marker gene coverage >= 3. Gene content was estimated by comparing the coverage from reads mapping to the MIDAS pangenome set to the single-copy marker gene set, and were considered present in our samples if having coverage greater than 1/3 or less than 3x the mean single-copy marker gene converge. SNV content was determined by mapping reads to the reference genome for the species, and sites were similarly masked on an individual basis if the coverage at the site was less than 1/3x or greater than 3x the genome-wide median coverage, To implement the quasi-phasing procedure (described below), we additionally require sites to have coverage >= 20 (Garud et al. 2019).

##### d) Quasi-phasing

To phase individual strain haplotypes, we follow the previously-described “quasi-phasing” (QP) procedure (Garud et al. 2019). In brief, samples harboring a dominant strain at high frequency for a given species can be confidently ‘quasi-phased’ by assigning high-frequency alleles as belonging to that dominant strain. Other alleles at intermediate frequency are denoted as missing from the haplotype. We use the same thresholds as in Garud et al. 2019, thereby using the same set of quasi-phaseable strains in Garud et al. 2019 and subsequent studies (Wolff and Garud 2024; Garud et al. 2019; Liu and Good 2024).

**e) Determining core genes**

We define ‘core genes’ as described in Garud et al. 2019. In brief, core genes were any genes present in at least 90% of QP strains for the species, where gene presence is defined as above in ‘Metagenomic processing with MIDAS’. We additionally exclude any genes with coverage >= 3 in any sample, as these may be genes promiscuously mapping reads from orthologous regions. Since the definition of which genes are core depends on the samples under consideration, we first infer a set of core genes among all quasi-phaseable samples. We then apply population-structure correction to these data (S3 Text). Subsequently, we re-calculate the set of genes considered core to the cluster of strains chosen post-population structure correction.

##### 3. Population structure correction

We first control for clonally-descended, closely related strains with an identical background across the much of the genome, which may artificially inflate haplotype sharing. To do so, only one strain among a cluster of closely-related strains with divergence in core genes < 5e-4/bp was retained in our analysis (Garud et al. 2019).

We then control for deeply-diverged subpopulations in gut commensal species with restricted gene flow (Truong et al. 2017; Costea et al. 2017), which can result in confounding signatures of elevated haplotype homozygosity. However, within subpopulations, lineages experience rapid decay in linkage disequilibrium (LD) due to high rates of gene flow (Garud et al. 2019; Liu and Good 2024). To identify such subpopulations, we extend the approach of Garud et al. 2019. First, we performed UPGMA clustering. By visual inspection, there are deep-level divisions between groups of strains. Garud et al. 2019 then chose the largest cluster representing a deeply diverged 'clade' to proceed in the analysis. However, with this approach we occasionally noticed runs of contiguous windows of H12 which all had the same value, suggesting incomplete correction, as there may exist clusters of strains with intermediate levels of population structure which are less visually apparent. Starting with every cluster of strains with at least 20 strains, we explicitly check LD decay, and then add in the next closely diverged strain iteratively (i.e. following the UPGMA algorithm), eventually testing LD decay among all clusters formed which contain 20 or more strains.

We use the following formula for LD:

$$r^2 = \frac{(pAB - pApB)^2}{pA\,(1 - pA)pB\,(1 - pB)} \qquad (3)$$

Where pA is the frequency of allele A; pB, the frequency of allele B, and pAB, the frequency of haplotypes carrying both A and B. We calculate $r^2$ for all 4D synonymous sites in core genes in two sets of sites: all sites up to 100 bp apart, and all sites 900-1100 bp apart. With sufficient recombination between alleles, average LD should decay precipitously between these two distances. In accordance with this expectation, LD decays within most clusters of samples, suggesting gene flow among lineages within clusters. However, when rates of LD decay (as measured by percent change in $r^2$) sharply drop, this indicates reduced gene flow, and often coincides with a noticeable increase in average genetic distance between strains. These represent clusters with strong population structure.

We thus define the population of strains for further analyses as the collection of lineages with highest percent decay in LD and lowest overall LD. After applying the correction, many species' population-corrected samples were composed of the same strains as in Garud et al. 2019. However 6 of 35 species differed. This additional correction improved the rate of LD decay within some species (Fig S3) and visually improved some scans by removing the runs of H12 mentioned above.

#### 4. Overview of bacterial simulations

We simulate recombinant bacterial populations using the forward-time simulator SLiM v. 4.0 (Haller and Messer 2019), with scripts inspired by (Cury et al. 2022). Since it is difficult to simulate realistic bacterial population sizes using forward time simulators, we scale all diversity and recombination parameters to a Wright-Fisher population of size $N_{sim}=10^4$ (Hoggart et al. 2007). We simulate chromosomes of length 100,000 bp. Crucially, we parameterize levels of nucleotide diversity and recombination to match individual gut commensal species, setting mutation rate $\mu = \frac{\bar{d}}{2 \cdot N_{sim}}$, where $d$ is the average nucleotide divergence at four-fold degenerate synonymous sites in core genes for that species.

In our simulations, we assume every individual has an equal chance of homologous recombination with other individuals. Reproduction is clonal, but homologous recombination occurs uniformly, with transfers from one individual to another characterized by a recombination rate $r$ (probability of recombination / bp / generation) and exponentially distributed tract length with mean $l_r$. To avoid recombination edge effects and better reflect bacterial populations, the chromosome is also modeled as being circular (where a recombination event can wrap around to the beginning of the chromosome), following the approach of (Cury et al. 2022).

To parameterize $l_r$, we set each species' mean tract length as the 0.63 quantile of observed tract lengths from Fig 3C in (Liu and Good 2024). This quantile associates with the mean of an exponential distribution. These tract lengths were inferred by the authors based on sequence divergence between closely related lineages, where one lineage obtained a recombinant fragment on an otherwise clonal backbone shared between the two strains. Rather than using the estimate of tract length from section S5 Text: "Determining the window size for H12 scans", we opted to use a length estimated independently of the LD decay curve, as we later compare the empirical LD decay curve to the simulated curve as a criteria of whether our simulations can recapitulate the empirical data, described below.

We set $r$ using species-specific apparent rates of recombination from Fig 1 of Liu and Good (2024). This rate comes from fitting a model of recombination to data from pairs of strains ranging from clonally-descended to highly mosaic. While not the only estimate of recombination rate from the work, this estimate may better reflect the "effective" recombination rate over long time scales of diversification and selection among the typically-diverged strains included in H12 scans.

Since $r$ cannot be directly observed from apparent rates of recombination (e.g. $r\,/\,m$, $r/\mu$), we convert Liu and Good's estimate from Fig 1 to $r$ using the following expression

$$r = \frac{r}{\mu} \cdot \mu = \frac{T_{mrca}}{T_{mosaic}} \cdot \frac{1}{\bar{d} \cdot l_r} \cdot \mu = \frac{l}{(1/\alpha - 1) \cdot l_r} \cdot \mu \tag{4}$$

Where α is obtained from the $T_{mrca}$ / $T_{mosaic}$ estimate from Fig 1 in Liu, $l_r$ is obtained as explained above, and μ is the simulation parameter as defined above.

We run each simulation for $10*N_{sim} = 10^5$ generations to establish a 'burn in', where the population has equilibrated to its stationary state. In each simulation, we sample the number of haplotypes that match the sample size for the species. If simulating hard sweeps, the beneficial mutation is seeded in the center of the chromosome after the burn in, and if simulating soft sweeps, the adaptive mutation rate $\Theta_A$ is turned on with a rate of 1.0 in the center of the chromosome after the burn in.

**a) Assessing the fit of simulations to data**

We ensure that the simulations recapitulate summary statistics in the data. While the simulations were constructed to match expected values of nucleotide diversity and recombination from estimates measured from the data, only those models that recapitulate multiple summary statistics in the data, including distributions of H12 and rates of decay in LD, can establish a credible neutral baseline for expected patterns of haplotype homozygosity in the absence of selection (Garud et al. 2021).

We first assessed whether observed haplotype homozygosity matches simulations. A match is deemed when the bulk of the simulated distribution (e.g. points lying between the 25th and 75th quantiles) overlap the bulk of the empirical distribution (e.g. Fig S2). In addition, we ensured that the empirical and simulated decay of LD over 10kb are visually similar.

We omitted the species *A. onderonkii*, *B. stercoris, B. intestinihominis, D. invisus* for lacking baseline distributions of H12 with low variance in the bulk. Of the 16 remaining species, 10 species visually recapitulated the distribution of H12 and rate of LD decay. See Table S2 for parameters used to simulate these species. We proceeded with these 10 species in calling significant H12 peaks (see S5 Text) and for all further analyses.

#### 5. Application of H12 to data

##### a) Determining the window size for H12 scans

H12 is calculated in genomic windows defined by a fixed number of variants. In that window, a haplotype is defined as a unique set of variants carried on a genomic background, and the haplotype's frequency is defined as the proportion of samples bearing the same haplotype. We identified species-specific window sizes in which local signals of excess homozygosity due to selection can be distinguished from the genomic baseline. Sufficiently large windows are unlikely to produce elevated haplotype homozygosity by chance. However, windows too large would likely preclude even the strongest sweeps from generating elevated haplotype homozygosity.

As H12 window sizes are defined as a fixed number of segregating sites, we chose a window size that, on average, spans a length by which LD is decayed (and H12 should be low), but is still sensitive to elevated LD from selective sweeps. As the rate of decay depends on the rate of recombination and the size of homologous tract lengths exchanged, we inferred a mean tract length directly from the LD decay curve, as explained below. Note that this tract length, $l_r^{\mathrm{LD}}$, inferred from the LD decay curve, is distinct from the Liu and Good tract length $l_r$ described previously, which the authors inferred from pairs of lineages exhibiting tracts of sequence divergence rather than from LD decay (Liu and Good 2024). In practice, we find that $l_r^{\mathrm{LD}}$ and $l_r$ have good concordance with each other (Fig S4). Using the estimate of the mean tract length $l_r^{\mathrm{LD}}$, we ran H12 scans using a window size defined as the average number of SNPs spanning 1x to 5x the tract length, ensuring a low baseline H12 value such that outlier windows corresponding to sweeps could be easily identified. See Table S1 for the window size in terms of SNPs and the corresponding genomic length.

We performed H12 scans on all species with at least 40 quasi-phaseable haplotypes passing population structure correction (S3 Text), including all contigs in which H12 could be computed in at least 50 windows.

**Inferring the mean tract length $l_r^{\mathrm{LD}}$**

We estimate the mean tract length, $l_r^{\mathrm{LD}}$, following methodology from (Wolff and Garud 2026). First, we fit the empirical decay of LD (as measured by $\sigma^2$) to the expected LD in a finite population with neutral mutation and drift (Ohta and Kimura 1969):

$$E(\sigma^2) = C\frac{10 + 2\,N\,R(l)}{22 + 26\,N\,R(l) + 4\,[N\,R(l)]^2} \tag{5}$$

Where C is an arbitrary normalization constant, and N is the effective population size. The recombination rate *R(l)*, represents the rate that two loci recombine at distance *l*. Under exponentially-distributed tract lengths:

$$R(l) = r l_r\left(1 - e^{-\frac{l}{l_r^{\mathrm{LD}}}}\right) \tag{6}$$

Where $r$ is the per-basepair recombination rate and $l_r^{\mathrm{LD}}$ is the mean tract length. As $\sigma^2$ should very closely approximate the linkage disequilibrium statistic $r^2$ at intermediate allele frequencies (McVean 2002), we calculated the empirical $r^2$ curve using all synonymous sites with MAF > 0.1 (calculated in 100 bp bins from 100 bp to 50 kb; see equation 3 for definition of $r^2$). We then fit these data to equation 4 to estimate $l_r^{\mathrm{LD}}$ using non-linear least squares.

**b) Control for missing data**

H12 is intended to be run on phased haplotype data in which missing data arises infrequently and randomly. However, metagenomic data may have high rates of missing data particularly in accessory genes (gene prevalence <0.9) resulting from strains lacking the accessory element. To account for this, in any H12 window, we only include a haplotype if it contains at least 70% of the window's variants (in practice, we did not notice drastic differences in scans when varying this threshold). We then cluster genomes that share the same haplotype if they exhibit no differences at non-missing sites. This occasionally results in a situation where a haplotype could join multiple haplotype clusters, even though those clusters exhibit SNV differences with one another. In this case, we randomly assign the haplotype to one of the clusters.

As this process of excluding haplotypes with large amounts of missing data will result in different numbers of haplotypes in different analysis windows, we standardize the number of haplotypes per window to a predefined sample size (Fig S1). For most species, we downsample to 80% of the total number of quasi-phaseable haplotypes. Using this approach, we were able to compute H12 for many contiguous windows for most species, with some small number of regions omitted. However, in the species *B. stercoris*, *B. thetaiotamicron, B. uniformis*, and *B. vulgatus*, we observed contiguous gaps of H12 windows, potentially due to the high levels of pan-genome content in the *Bacteroides* (Zou et al. 2019), making it hard to determine a baseline value for H12. To be able to analyze these species, we lowered the downsample proportion to 50% the number of quasi-phaseable haplotypes. See Table S1 for downsample numbers for individual species.

**c) H12 computed on synonymous sites carries greater power to detect sweeps**

H12 has typically been calculated using any SNP, whether synonymous, non-synonymous, or non-coding (Garud et al. 2015). In this paper, we calculate H12 in windows composed exclusively of synonymous sites (not just 4D sites, but any synonymous site). Unlike eukaryotic genomes, the vast majority of sites in commensal gut bacteria are coding sequences, implying that windows composed of synonymous sites can still cover the majority of the genome and not have an excessively variable distribution of window lengths.

In simulations, we found that calculating H12 in windows composed of synonymous sites results in increased power to detect selective sweeps compared to H12 windows composed of synonymous and non-synonymous variants (Fig S5). Our simulations consisted of both hard and soft ($\Theta_A = 1$) selective sweeps across the following beneficial selective coefficients ($s_b \in$ 0.01,0.03,0.05,0.1), conducting 200 simulations per sweep type and selective coefficient combination. Nonsynonymous sites were modeled as having selective coefficient -0.001, and comprised 2/3 of *de novo* mutations, while the remaining third were modeled as synonymous, and neutral. Other simulation details match S4 Text ("Overview of bacterial simulations"), with $\bar{d}$ set to 0.005 and $r/\mu$ set to 0.05. The average tract length exchanged was set to 10kb, reflecting the typical tract length observed in data. To compare the ability of H12 calculated on only synonymous sites vs all sites, we first parsed exclusively synonymous variants from all simulations. So that the density of all segregating sites was matched to the density of synonymous sites, we separately downsampled all variants in a simulation to the number of synonymous variants in that simulation. We then calculated H12 on both parsed sets of haplotypes.

Understanding why H12 has increased power when computed on synonymous variants only is an area for future work. However, one reason excluding nonsynonymous variants may help is that bacteria are gene-dense, resulting in a high number of nonsynonymous variants, which are additionally under purifying selection. The presence of many rare deleterious variants may reduce haplotype homozygosity, making it more difficult to detect a sweep.

**d) Removal of regions with low recombination rate**

A decline in local recombination rate may spuriously lead to high haplotype homozygosity (O'Reilly et al. 2008). Previous work has shown variation in homologous recombination rates in gut commensals across the genome (Liu and Good 2024). To confirm that H12 outliers are not confounded by regions of low recombination, we implemented an additional control by calculating the apparent recombination rate of observed recombination events in gut commensal species in 50kb windows using recombination events inferred in a previous work (Liu and Good 2024). Any windows with a recombination rate < 20% the average genome-wide recombination rate are plotted in grey dots in scans (Fig 2F, Fig S6), and windows in these regions were excluded in future analyses.

**e) Calling significant H12 peaks**

To identify putative sweeps in *C. difficile* and *N. gonorrhoeae*, we highlight all H12 values exceeding the 99th percentile of the genome-wide distribution of H12 values as the parameters required to simulate neutral evolution were not available for these species. For gut commensal species, we instead assess whether H12 values are significantly greater than expected under simulations of neutrality using a critical value tuned to a 1-per-genome false positive rate. This critical value is derived from the distribution of H12 values measured from simulations of panmictic, neutrally evolving haploid populations with rates of nucleotide diversity and recombination matching empirical estimates for each species as described above in S4 Text (Fig S2).

The significance threshold was calculated on a species-by-species basis, from 10,000 simulated independent neutral estimates of H12. The 1-per-genome false positive threshold was defined to be the *n*th highest neutral H12 value, where *n* is 10,000 / the number non-overlapping windows in the scan, rounded up. We use the number of non-overlapping windows because each of the simulated neutral values is independent, but even non-overlapping windows display positive dependency, making the threshold conservative with respect to false positives (Benjamini and Yekutieli 2001). Across our scans, *n* was at minimum 17, indicating our 1-per-genome critical value is robustly sampled for every species.

To identify individual sweep events, we called 'peaks' by clustering all windows containing at least 10 consecutive windows above the significance threshold. The peaks were called one-by-one, starting with the cluster containing the highest H12 value, and omitting any clusters lying within two average window lengths (in terms of kb) from further peak assignment. Then the second-highest H12 cluster is assigned, and so on, until no clusters remain.

In the species *Bacteroides thetaiotamicron* and *Alistipes putredinis*, the significance threshold was low enough that a substantial portion of windows exceeded it. For downstream analysis, we set a more conservative significance threshold of the 99% empirical H12 quantile, resulting in 2 peaks for *B. thetaiotamicron* and 4 peaks for *A. putredinis*.

#### 6. ABC classification of soft and hard sweeps

##### a) Overview

We use approximate Bayesian computation (ABC) to quantify support for hard vs soft sweeps for each sweep, following (Garud et al. 2015). Specifically, we use Bayes' factors to compare the likelihood that the haplotype structure at a sweep was generated under a hard vs. soft sweep model. We summarize each peak's haplotype spectra using three summary statistics: H12 (defined in the main text), H2/H1, and hamming12 (defined below).

For each species, we simulate 300,000 hard and 300,000 soft selective sweeps, drawing the parameters for beneficial selection ($s_b$), partial frequency (PF), and ending time of sweep ($T_e$) as follows: $s_b \sim \text{Unif}(0,0.2)$, PF ~ Unif(0,1) and $T_e \sim \text{Unif}\{0, 200\}$ (i.e. discrete whole numbers). Soft sweeps were simulated with $\Theta_A$ of 1.0, which should generate almost exclusively soft sweeps (Pennings and Hermisson 2006) and hard sweeps were simulated by seeding a single beneficial mutation.

We can then compare empirical values of H12, H2/H1, and hamming12 at inferred sweeps to those calculated from simulations. The posterior probability of the soft sweep and hard sweep model is proportional to the number of soft sweep and hard sweep simulations matching the empirical sweep, respectively. Specifically, we determine the strength of the evidence for soft vs. hard sweeps by calculating the Bayes' Factor between the following models:

$M_h$ := Model where peak is hard
$M_s$ := Model where peak is soft under $\Theta_A = 1.0$

We calculated Bayes factors by taking the ratio of the number of matching simulated soft sweeps to matching simulated hard sweeps. In practice, as no simulation can exactly match empirical values of H12, H2/H1, and hamming12 values, we include all simulations within a Euclidean distance < 0.25 from the observed values $H12_{obs}$, $H2_{obs}/H1_{obs}$ and Hamming12 $_{obs}$ for each set of

300,000 soft versus hard sweeps. We calculated the Euclidean distance between simulation i and the observed data as follows:

$$d_i = \sqrt{\left(\frac{\mathrm{H12_{obs}} - \mathrm{H12_i}}{\mathrm{sd(H12)}}\right)^2 + \left(\frac{\mathrm{H2/H1_{obs}} - \mathrm{H2/H1_i}}{\mathrm{sd(H2/H1)}}\right)^2 + \left(\frac{\mathrm{hamming12_{obs}} - \mathrm{hamming12_i}}{\mathrm{sd(hamming12)}}\right)^2} \quad (7)$$

Here, *sd* refers to the standard deviation of the summary statistic amongst all simulations of that model (soft or hard).

**b) Summary statistics of haplotype structure**

**H2/H1**

To distinguish hard from soft sweeps, we use two summary statistics for ABC in addition to H12. The first statistic is H2/H1 (Garud et al. 2015), where

$$H1 = \sum_{i \geq 1}^{n} p_i^2 \quad (8)$$

and

$$H2 = \sum_{i \geq 2}^{n} p_i^2 = H1 - p_1^2 \quad (9)$$

H1 is haplotype homozygosity and H2 is haplotype homozygosity excluding the contribution of the most common haplotype. H2/H1 describes the proportion of homozygosity without the most common haplotype and is expected be large for a soft sweep and small for a hard sweep when H12 is elevated.

**Hamming12**

Haplotype frequencies alone can differentiate between hard and soft sweeps in the majority of cases. However, in a hard sweep the sweeping haplotype can be occasionally broken up by mutation or recombination events over the course of the sweep, resulting in multiple haplotypes at high frequency, albeit with low divergence from one another (Hermisson and Pennings 2017). While the distribution of haplotype frequencies in this 'broken hard sweep' scenario may resemble a soft sweep, a 'broken hard sweep' contrasts with the expectations of a soft sweep from recurrent mutation because, in the latter case, independent adaptations should be seeded on random haplotypes from the population; thus haplotypes at high frequency for a soft sweep should exhibit higher divergence from one another than a broken hard sweep.

To better distinguish between hard and soft sweeps, we introduce an additional summary statistic, hamming12, which measures the hamming distance between the first- and second-most frequent haplotypes, normalized by the typical distance between two randomly chosen haplotypes of the same length. Most hard sweeps (not broken), as well as soft sweeps from *de novo* mutations, are expected to have hamming12 values of 1, consistent with choosing any two random haplotypes. In this case, haplotype statistics (e.g. H2/H1) best distinguish hard from soft sweeps. However, in a 'broken hard sweep' scenario, hamming12 should be low, making hard sweeps distinct from soft sweeps even when haplotype frequencies alone are uninformative. However, we note it is also possible for a soft sweep to arise from standing genetic variation (SGV), whereby a mutation has time to segregate in the population and arise on multiple haplotype backgrounds before the onset of selection. In this case, the sweeping haplotypes bearing the adaptive allele will be more closely related to each other than two random haplotypes drawn from the broader population. However, they will not be as closely related as in the case of a broken hard sweep. We find that soft sweeps from SGV have higher and distinct values of H2/H1 compared to hard sweeps, and are still primarily classified as soft (Fig S8). In total, we show hard sweeps become more distinguishable from soft sweeps when all three statistics (H12, H2/H1, hamming12) are used (Fig S10).

#### 7. Stochastic model of a sweep across host microbiomes

##### a) Overview

We consider the population genetic processes that might allow for sweeps to spread across a metapopulation of human gut microbiomes. However, given the intractability of a metapopulation model of millions of bacterial populations, we make a number of conservative assumptions that allow for a higher probability of observing hard sweeps. With these assumptions, we arrive at the well-characterized Chinese restaurant process, from which we can generate predictions representing the upper bound of observing hard sweeps arising from a given mutation and recombination rate. By biasing our model to hard sweeps, that makes it all the more surprising that neither hard nor soft sweeps are observable in the empirical range of these rates, implying that widespread parallelism should be predominant in this regime. We list these assumptions below, describe the resulting stochastic process, and provide the analytical expressions from which we determine the number of *de novo* adaptive origins and frequencies of origins after the selective sweep has finished.

##### b) Model assumptions

1. We assume each human gut microbiome harbors only one strain. In reality hosts are oligocolonized from with ~1 – 5 divergent strains (Garud et al. 2019; Truong et al. 2017). However, this simplification ensures that at most one novel haplotype can arise from a *de novo* adaptive mutation event within the host.

2. We assume only one human gut microbiome experiences a selection pressure at a time. This is a conservative assumption with respect to observing hard sweeps because every previously adapted gut microbiome is now available to contribute an adaptive origin via recombination to the microbiome currently under selection. The opposite extreme, where the entire host metapopulation synchronously shares the selective pressure, requires

sweeps spreading across hosts to outpace the time to fixation of a *de novo* mutation within individual hosts, which is unlikely.

3. We assume the fixation time of the within-host sweep, once seeded with an adaptation, to be instantaneous relative to the time of spread of the across-host sweep. In other words, the next host population does not adapt until the adaptive allele has swept to fixation within the prior host. With a similar logic to 2., this increases the likelihood of a given gut microbiome acquiring an allele via recombination, thereby increasing the probability of hard sweeps.

4. We constrain the number of adaptive origins per host to be 1. That is, the microbiome currently under selection can only acquire an adaptive mutation via *de novo* mutation or recombination from one other microbiome since we assume that the mutation instantaneously fixes within the host that acquires the mutation.

5. As we only consider the dynamics of a single locus, we do not explicitly model the process of strain replacement. However, as the rate of strain replacement is much lower that of recombination (or *de novo* mutation) (Garud et al. 2019), this process can be approximated by very slightly increasing the rate of recombination relative to mutation, which does not affect the qualitative outcomes of our model.

6. Finally, we assume each host's strain may freely recombine with any other host's strain. Previous work has established that population structure increases the likelihood of soft sweeps (Pennings and Hermisson 2006; Feder et al. 2019; Ralph and Coop 2010; Paulose et al. 2019). Therefore, this assumption is conservative by decreasing the number of origins and increasing the probability of hard sweeps.

**c) Modeling the spread of an adaptation with the Chinese restaurant process**

Under the above assumptions, we consider the probability that an adaptation in a given host will be acquired via *de novo* mutation vs a recombination event from a different host already harboring the adaptation. By the superposition property of point processes, the identity of the adaptive origin within a host is determined by a single categorical draw, with probabilities parameterized by the competing rates of *de novo* mutation vs recombination from other hosts currently carrying the adaptation.

Consider an across-host sweep of 2 hosts, with host 1 already containing an adaptive origin. Host 2 can then obtain the adaptive fragment from host 1 or *de novo* mutate a new origin. We can consider this scenario as partitions of the set {1, 2}, where sharing a partition denotes sharing an origin: {{1, 2}} (recombination) vs. {{1},{2}} (mutation). We parameterize these events as follows:

$$P(\text{adaptation via recombination / generation}) = \mathrm{r} * \mathrm{l_r}$$

$$P(\text{adaptation via mutation / generation}) = \mu$$

$$\Rightarrow \mathrm{P}(\{\{1,2\}\}) = \frac{r * l_r}{\mu + r * l_r}$$

$$\Rightarrow \mathrm{P}(\{\{1\},\{2\}\}) = \frac{\mu}{\mu + r * l_r}$$

Where $\mu$ is the mutation rate per bp per generation, $r$ is the recombination rate per-bp per generation (between the two strains, in this case), and $l_r$ is the average tract length exchanged in a recombination event.

We now extend this to an across-host population of size N hosts. A host obtains a particular origin $k$ via recombination at a rate proportional to the number of hosts already containing that origin, $n_k$:

$$\mathrm{P}(\text{adaptation from } k) = \mathrm{r} * l_r * \frac{n_k}{N} \quad (10)$$

The first host to adapt, trivially, must mutate a *de novo* origin. With these assumptions, and viewing the across-host sweep process as a taking values of the set of partitions of all possible ways to partition the set {1, …, N} (Yee Whye Teh 2017), our stochastic model describes a Chinese restaurant process with parameter $\theta = \frac{N\,\mu}{r\,l_r}$. We use this result to find the expectation for when N is $10^8$, approaching census human population sizes in the industrialized world, but not requiring the adaptation to be universally beneficial, as smaller N is more conservative with respect to reducing the number of origins and observing hard sweeps.

The odds of host *i* acquiring an adaptation via recombination vs. mutating a new origin is then

$$\frac{P(adaptation\ via\ recombination)}{P(adaptation\ via\ de\ novo\ mutation)} = \frac{r * l_r * f_A}{\mu} \tag{11}$$

Where the adaptive across-host allele frequency $f_A = \frac{i-1}{N}$ because every previous i - 1 hosts have either mutated or acquired an origin. One way of interpreting this ratio is its resemblance to the bacterial recombination metric *r/m*, which describes the rate at which a site diversifies via recombination vs. mutation:

$$\frac{r}{m} = \frac{r * l_r * \bar{d}}{\mu} \tag{12}$$

Where $d$ is nucleotide diversity at the site. However, in equation 11, the adaptive allele frequency $f_A$ is substituted. Thus, $\frac{r * l_r * f_A}{\mu}$ describes the recombination-mutation ratio by which a given host obtains an adaptation. With $f_A$ a dynamic property of the stochastic process as the allele spreads across hosts, we evaluate the frequencies and numbers of origins across the parameter space of $\frac{r * l_r}{\mu}$, varying the relative contribution of mutation and recombination.

We set the population size N to be $10^8$ in the Fig 4. However, to test how sensitive our model is to different population sizes, we additionally vary N while fixing $\frac{r * l_r}{\mu}$ at 1000, a value typical of the empirical range of gut bacteria (Fig S12) (Liu and Good 2024). Even when varying N across several orders of magnitude, our model predicts that any adaptive origin would not achieve appreciable frequency in the population assuming $\frac{r * l_r}{\mu} = 1000$. Instead an orders of magnitude difference in $\frac{r * l_r}{\mu}$ would be required to observe soft or hard sweeps.

**d) The expected number of origins and origin frequencies under the Chinese restaurant process**

Rather than explicitly simulating the Chinese restaurant process under such large N, we instead use expressions of its expected behavior. The expected number of origins is the sum of the probabilities that each host establishes a new origin:

$$\mathrm{E}\left(N_{origins}\right) = \sum_{i=1}^{N} \frac{\mu}{\mu + \frac{i-1}{N} \cdot r \cdot l_r} = \sum_{i=1}^{N} \frac{\theta}{\theta + i - 1} = \theta \cdot \left(\Psi(\theta + N) - \Psi(\theta)\right) \quad (13)$$

Where $\theta = \frac{N\mu}{r\, l_r}$ and $\Psi(.)$ is the digamma function (Yee Whye Teh 2017).

The expected frequency of the *k*-th most frequent origin is the sum of the probabilities that each host harbors that origin. Denote $p_\beta(i, k)$ as the probability that host *i* harbors origin *k*, and $p_\gamma(i, k)$ as the probability there exists *k* origins at the time host *i* adapts. Then, the expected frequency of *k* is (passerby51 2021):

$$\mathrm{E}(f_k) = \frac{1}{N} \sum_{i=1}^{N} p_\beta(i, k)$$

$$= \frac{1}{N} \{p_\beta(1, k) + (N-2) \cdot p_\beta(2, k) + \sum_{i=2}^{N-1} \sum_{z=1}^{i} \frac{\theta}{\theta + z} [p_\gamma(z, k-1) - p_\gamma(z-1, k-1)]\} \quad (14)$$

Where $p_\beta(1, k) = \mathbf{1}\{k = 1\}$, and $p_\beta(2, k) = \frac{1}{\theta + 1}\mathbf{1}\{k = 1\} + \frac{\theta}{\theta + 1}\mathbf{1}\{k = 2\}$. The function $p_\gamma(i, k)$ is the probability mass function of the Chinese restaurant distribution (Yee Whye Teh 2017):

$$p_\gamma(i, k) = \frac{\Gamma(\theta)}{\Gamma(i + \theta)} \cdot |S(i, k)| \cdot \theta^k \quad (15)$$

Where S(i,k) are Stirling numbers of the first kind and $\Gamma(.)$ is the gamma function. We approximate Stirling numbers of the first kind for $i > 20$, $k < 10$ with the following expression:

$$\mathrm{S}(i, k) = \frac{\Gamma(i)}{\Gamma(k)} \cdot log(i)^{k-1} \quad (16)$$

In practice we found this approximation to have good concordance with a simulated Chinese restaurant process for N = 100,000 and θ = 100.

**e) Approximating the waiting time for adaptations to arise under reciprocal pairwise epistasis**

Our stochastic model shows that $\frac{r * l_r}{\mu}$ must be several orders of magnitude larger than empirically estimated to observe hard or soft selective sweeps. One way to achieve a larger value is if the adaptive mutation rate is much smaller than assumed for a single site. To illustrate why such a reduction in mutation rate may be plausible, consider the waiting time for an adaptation to arise. For a single site adaptation, the adaptive mutation rate is

$$\Theta_A \sim (N \cdot f) \cdot \mu \quad (17)$$

where N is the number of bacterial cells in a host, $f$ is the species relative abundance, and $\mu$ is the per-site mutation rate. With estimates of N of $10^{13}$ (Sender et al. 2016), $f \sim 10^{-2} - 10^{-1}$, and $\mu$ of

$10^{-10}$-$10^{-9}$ (Barrick and Lenski 2013; Sung et al. 2012) and assuming 1-10 generations/day (Korem et al. 2015; Ghosh and Good 2022), a single site adaptation should arise within an individual gut at least once per day, as previously estimated (Zhao et al. 2019).

Now suppose an adaptation requires reciprocal pairwise epistasis between two mutations that are strongly deleterious alone but beneficial together. The double mutant has a mutation rate of $\mu^2$, lowering $\Theta_A$ to ~$10^{-6}$ – $10^{-9}$ per generation. This would result in a waiting time for the double mutant to appear in any given host on the order of hundreds to millions of years. Thus, a double mutant adaptation is unlikely to ever arise in any given individual in their lifetime.

If selection pressures are shared across hosts, however, the double mutant can arise in any of these hosts. Now,

$$\Theta_{A\ (across\ hosts)} \sim \mathrm{N_{hosts}} \cdot (\mathrm{N} \cdot \mathrm{f}) \cdot \mu^2. \quad (18)$$

Assuming the adaptation is adaptive in any one of $10^8$-$10^9$ hosts, (e.g. spanning ~10% of the industrialized world to ~10% of the global population), $\Theta_{A\ (across\ hosts)}$ ~ 0.1 to 1000 per generation. In other words, the waiting time for the double mutant drops to a matter of days in a broader population of hosts. With this rate, the adaptation may be able to spread to high frequency across host microbiomes primarily via recombination, leading to observable sweeps. More generally, with the human population size being on the order of bacterial mutation rates, complex epistatic adaptations that have a near-zero likelihood of being generated within individual hosts can be readily attained across hosts. Furthermore, epistasis may be one of many types of complex adaptive architectures which have a low mutation rate. For example, structural or accessory elements in a bacterial genome likely cannot *de novo* mutate on the timescale of a sweep even in the largest of bacterial population sizes.

8. **Genomic signatures of complex adaptation**

**a) Calculating nucleotide diversity quantiles**

In Figure 5C, we compare the relative depression in nucleotide diversity at each of the 55 sweeps identified with H12 to expectations under hard sweeps, soft sweeps, and neutrality. For each sweep, we compute nucleotide diversity, $\pi$, in 1000 bp windows centered around each H12 window's central SNP and expressed it as a quantile of the genome-wide distribution. The genome-wide distributions of $\pi$ was calculated in sliding windows of 1000 bp with a step size of 100 bp, excluding any windows that had been excluded from the H12 scan.

We compare the distribution of quantiles observed in the data to simulated sweeps. To do so, we simulated 1000 hard sweeps and 1000 soft sweeps ($\Theta_A$ = 1.0), and calculated $\pi$ in 1000 bp windows around the beneficial allele. We then assessed the sweep's quantile of $\pi$ in 2000 neutral simulations (i.e. analogous to the corresponding genomic 'baseline'). The simulation parameters used were roughly matching mean values observed across species: tract length $l_r$ = 10kb, $r / \mu$ = 0.1, and $d$ = 0.01, sample size of 100. Selection strength, partial frequency, and time since end of the sweep were drawn from uniform distributions matching the priors described in S6 Text.

### Supplemental Figures

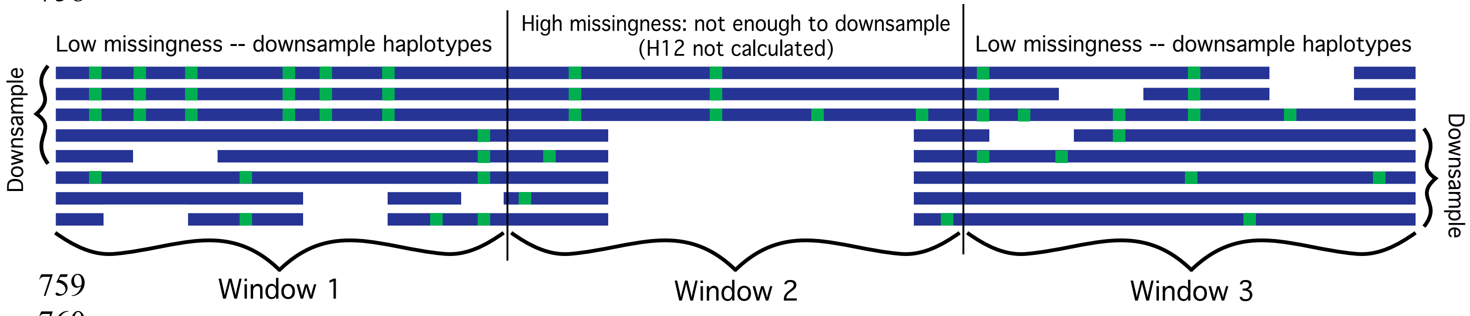

**Figure S1. Overview of H12 with downsampling.**
H12 is calculated in windows of a pre-defined number of variants. Any haplotype containing >= 30% missing data is removed from the window. Windows may be excluded for not having a minimum number of complete haplotypes (Window 2). Windows containing more than the minimum number of complete haplotypes are downsampled to allow comparison. In the figure above, the blue and green areas denote haplotypes where the major and minor alleles have been called, while the white areas denote missing data.

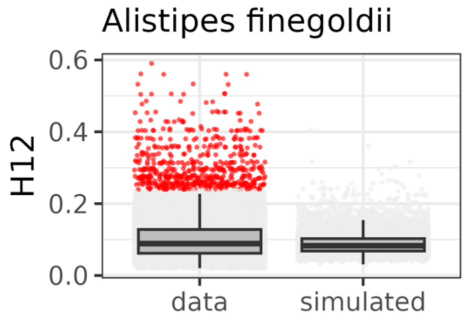

**Figure S2. Elevation of empirical distributions of H12 compared to neutral simulated distributions of H12 in *Alistipes finegoldii.***

We simulate a critical threshold beyond which H12 values are inconsistent with neutrality, tuned to a false positive level of 1-per-genome. Shown here is the distribution of empirical and simulated H12 in *A. finegoldii*. Windows passing the significance threshold (shown here in red) are clustered into peaks identifying individual sweep events.

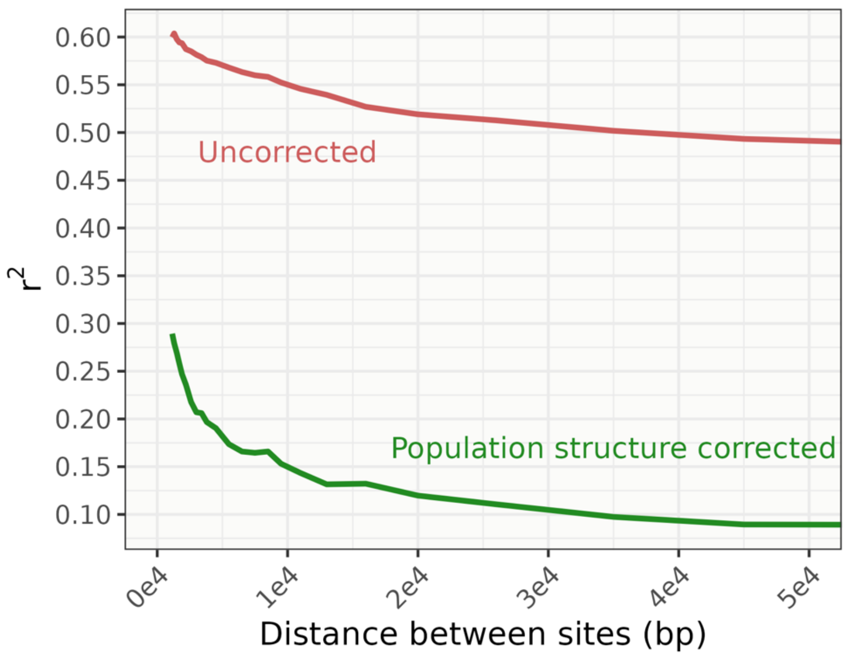

**Fig S3. Population structure correction improves the decay of linkage disequilibrium in *B. vulgatus***

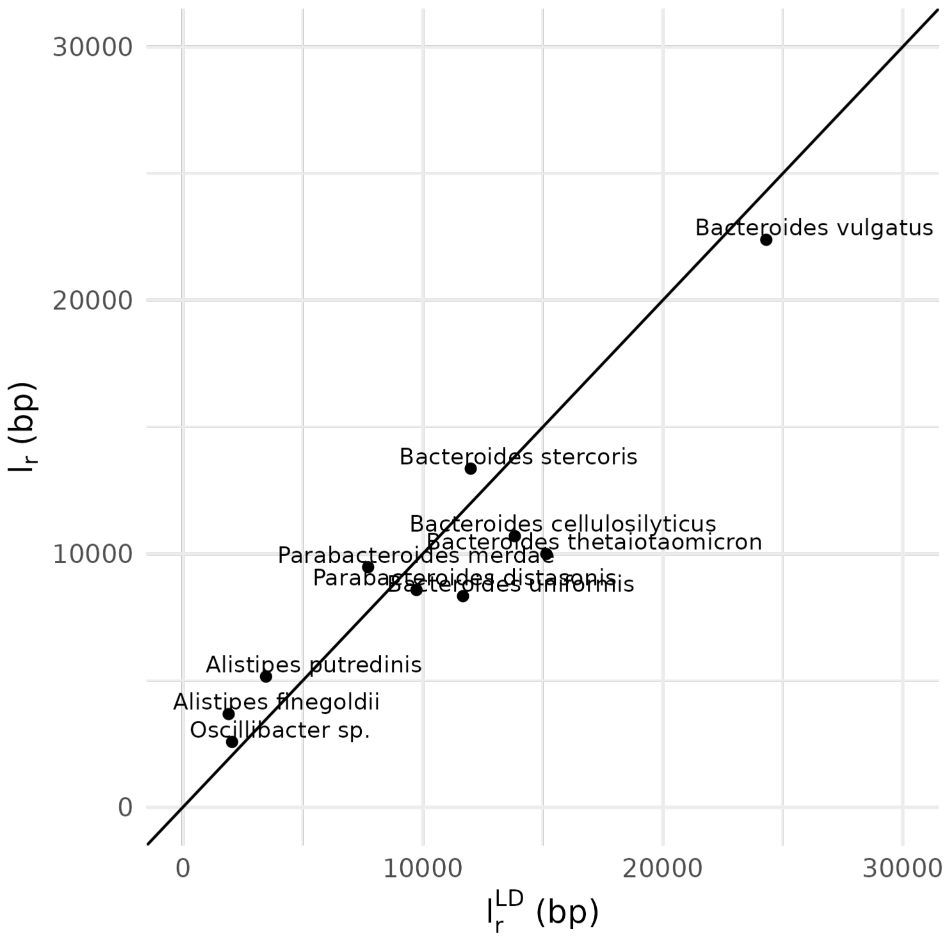

**Fig S4. Comparison of mean tract lengths $l_r$ and $l_r^{LD}$ among gut commensal species.** The estimate of $l_r$ used for simulations is derived from the empirical distribution of tract lengths inferred in (Liu and Good 2024). The estimate of $l_r^{LD}$ used to determine H12 window sizes for all commensal and pathogenic species is inferred directly from the decay of linkage disequilibrium (equations 5 and 6). Both estimates have high concordance among the 10 gut commensal species analyzed in this work.

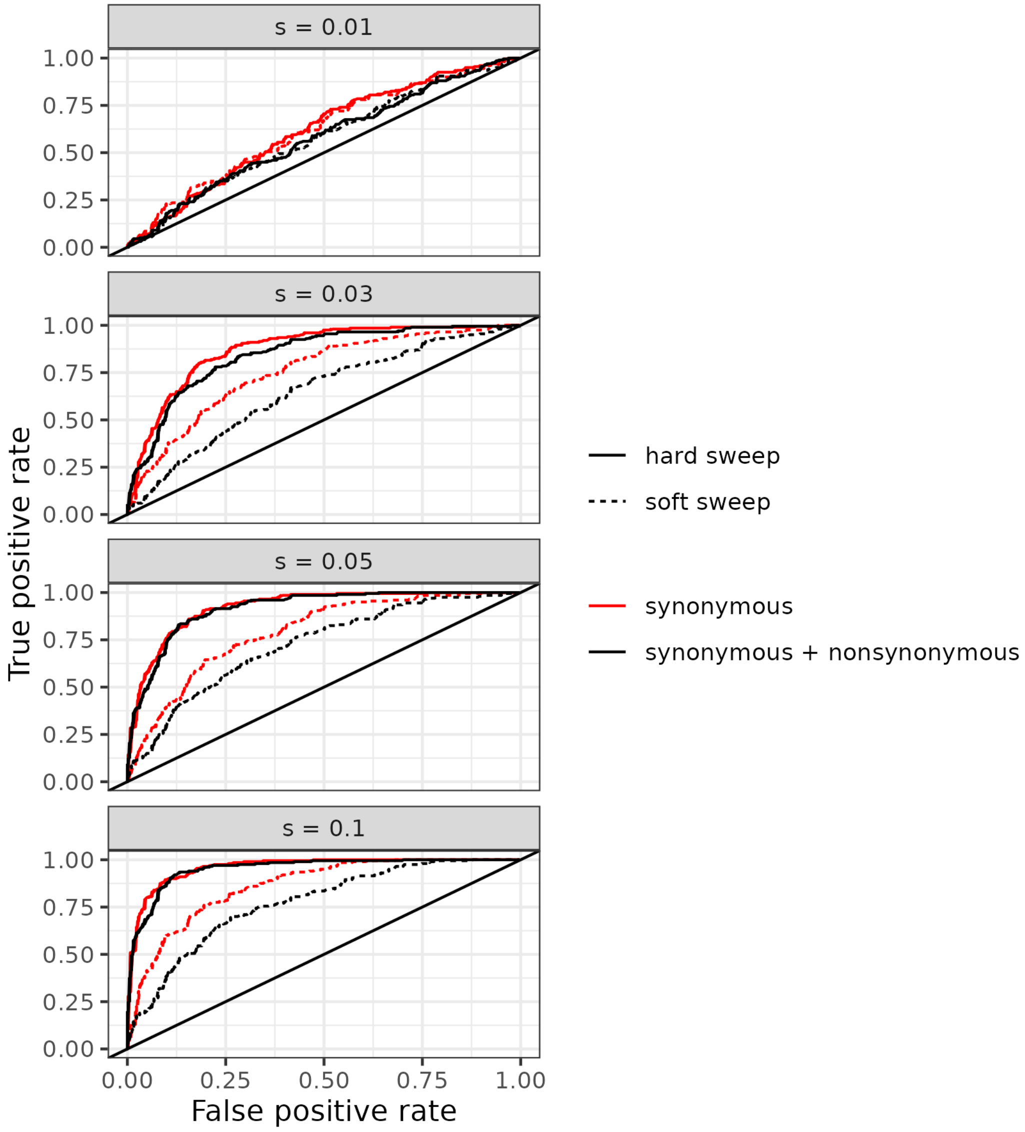

**Fig S5. H12 computed on synonymous variants has increased power when compared to H12 using synonymous and nonsynonymous variants.**
Both neutral (synonymous) and deleterious (nonsynonymous) variants were simulated with selection coefficients of 0 and $10^{-3}$, respectively, and Ne=$10^4$. Following a 10*Ne 'burn-in', a selective sweep with beneficial selection coefficient as specified in the figure was seeded. Hard sweeps were simulated by seeding a single beneficial mutation, and soft sweeps were simulated with an adaptive mutation rate, $\Theta_A$, of 1.0. ROCs were constructed from 200 simulations with the selective sweep and 200 without the selective sweep, for each sweep type and selective coefficient.

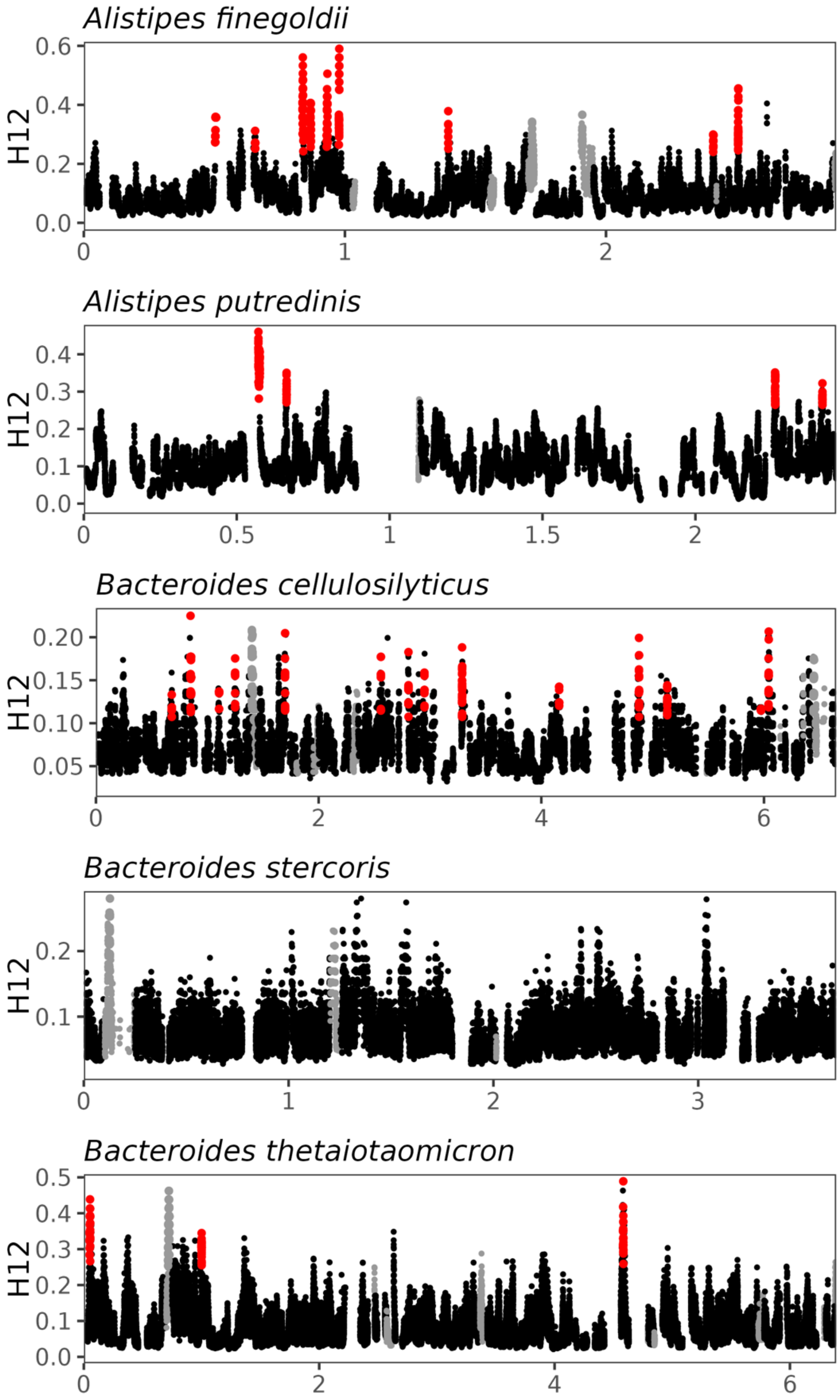
Alistipes finegoldii
H12
0.0
0.2
0.4
0.6
0
1
2
Alistipes putredinis
H12
0.0
0.1
0.2
0.3
0.4
0
0.5
1
1.5
2
Bacteroides cellulosilyticus
H12
0.05
0.10
0.15
0.20
0
2
4
6
Bacteroides stercoris
H12
0.1
0.2
0
1
2
3
Bacteroides thetaiotaomicron
H12
0.0
0.1
0.2
0.3
0.4
0.5
0
2
4
6

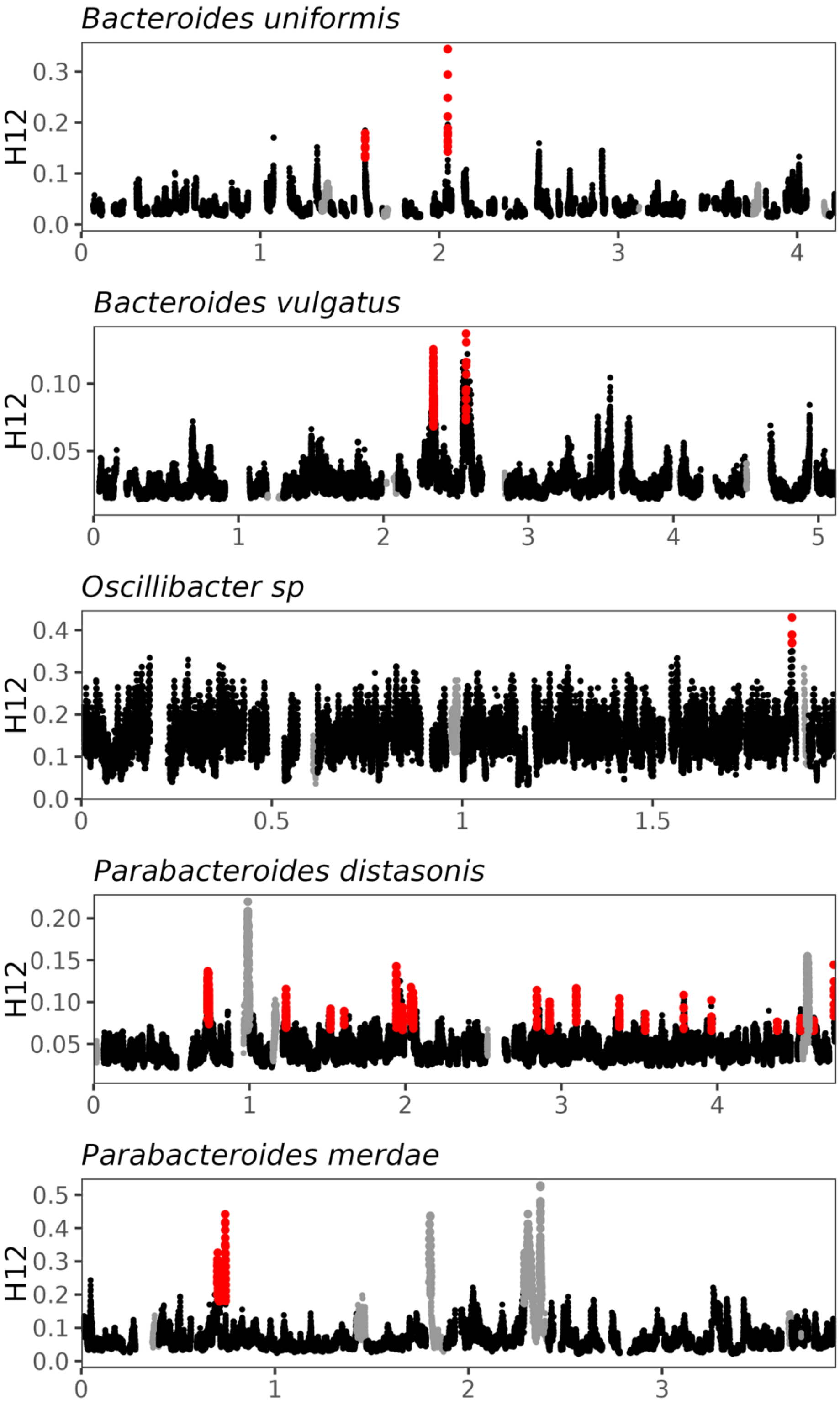

**Figure S6. H12 scans in gut commensal species**. Grey dots denote windows with low recombination rates, while red dots denote peaks of H12 values representing individual selective sweeps. The x axis for all scans denotes genomic position (Mb).

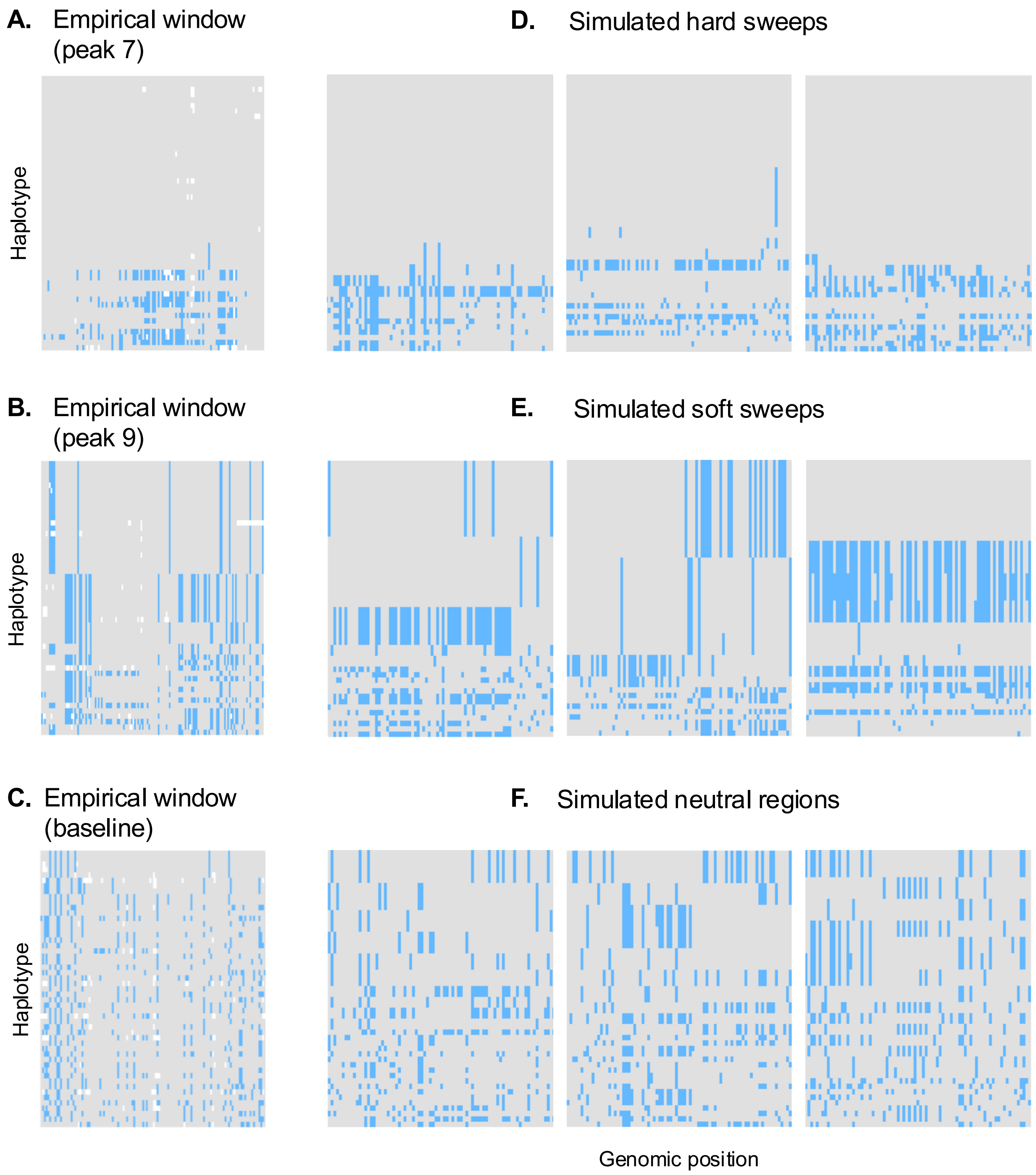

**Figure S7. Haplotype diversity in *A. finegoldii* at H12 peaks compared to simulated hard and soft selective sweeps**

On the left are haplotypes from H12 windows in *A. finegoldii*. On the right are haplotypes from simulations of (**D**) hard sweeps, (**E**) soft sweeps ($\Theta_A$ = 1.0), and (**F**) neutral regions with matching number of SNPs as the empirical window. Blue indicates minor allele.

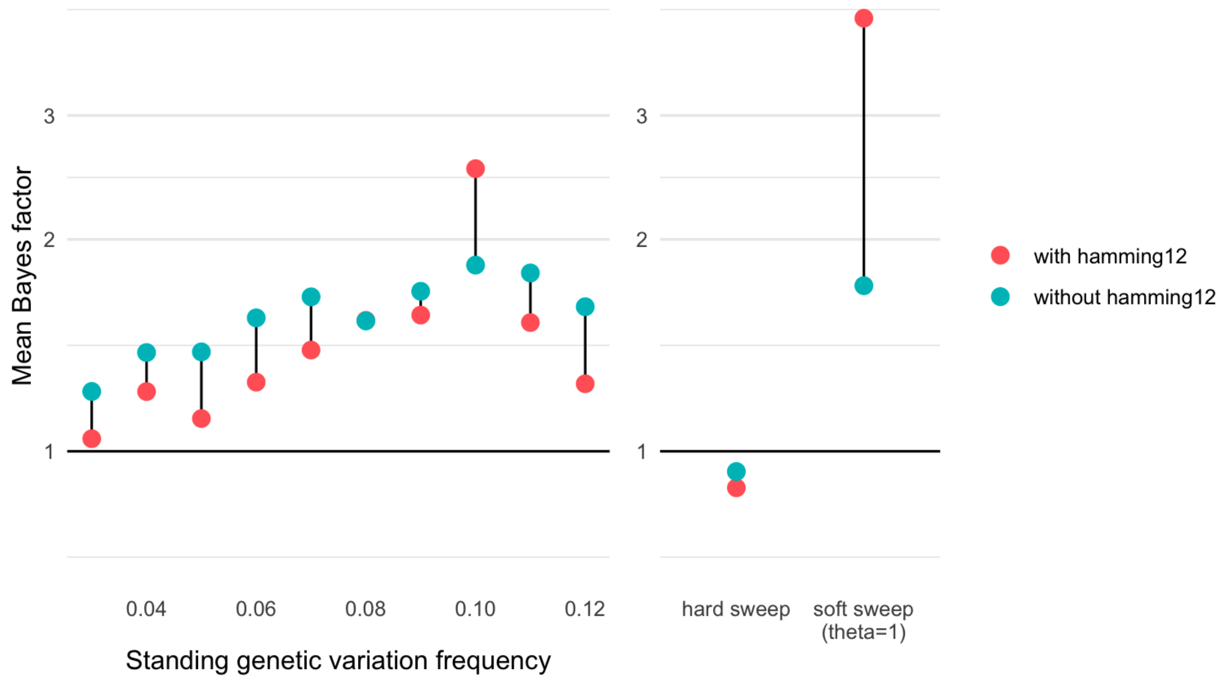

**Figure S8. Soft sweeps arising from SGV are still distinguishable from hard sweeps when using hamming12.** Simulations of hard and soft sweeps from standing genetic variation (SGV) were conducted using SLiM and matching ABC parameters for *A. finegoldii*. To simulate SGV sweeps, the adaptive mutations were allowed to segregate and drift to a predetermined frequency ({0.03, 0.04, …, 0.12}) before selection is enabled, with 200 simulations conducted for each frequency. Plotted are the mean Bayes' factor for SGV sweeps, when assessed on the *A. finegoldii* posterior for soft sweeps simulated from recurrent mutation. All SGV soft sweeps have a mean Bayes factor >=1, which is only slightly lowered when hamming12 is included as a summary statistic. Hard sweeps and soft sweeps from recurrent mutation are also plotted as a comparison, which both show improved classification with the inclusion of hamming12.

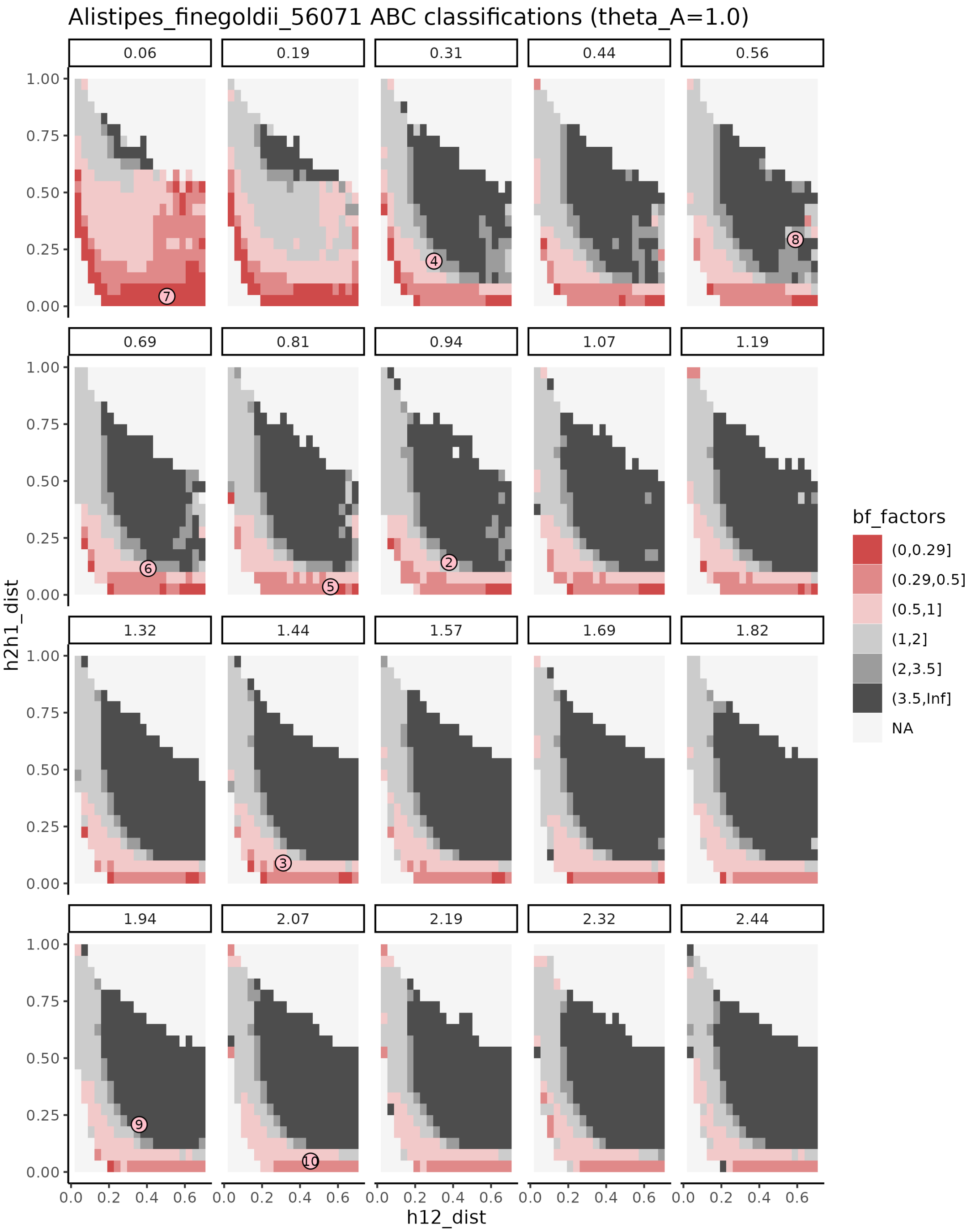

**Figure S9. Distribution of Bayes factors across hamming12, H12 and H2/H1 for *Alistipes finegoldii*.**
Faceting across different values of hamming12, and we plot the H2/H1 vs. H12 landscape of BF support for soft sweeps vs. hard sweeps. The darker grey denotes higher-confidence support for soft sweeps, while dark red shows high support for hard sweeps. Higher values of H2/H1 in general show greater support for soft sweeps, but the landscape is highly dependent on hamming12, with lower values of hamming12 providing greater support for hard sweeps across much of the joint distribution of H12 and H2/H1. Individual dots represent empirical sweeps, where the number corresponds to the peak's location, from left to right (Fig 2F).

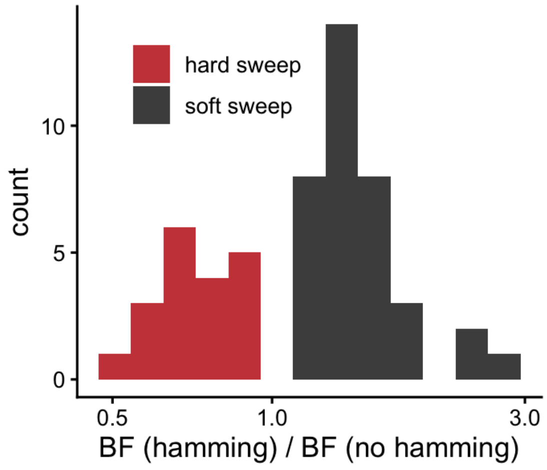

**Figure S10. hamming12 improves ABC classification at sweep posteriors**
For each sweep, we simulated 1000 sweeps from the soft or hard sweep posterior distribution, whichever matched the initial BF classification. These posterior predictive replicate sweeps were then reassessed on the original BF landscape, and the mean ratio of the BF assessed with hamming12 and without hamming12 is plotted for every sweep. The inclusion of hamming12 as a summary statistic improved the classification of every soft sweep (ratio of BF > 1), and improved the classification of every hard sweep (ratio of BF < 1), suggesting that hamming12 increases certainty by a factor of 2-3 in distinguishing between hard and soft sweep models for sweep replicates, and improves on summary statistics of haplotype frequencies alone.

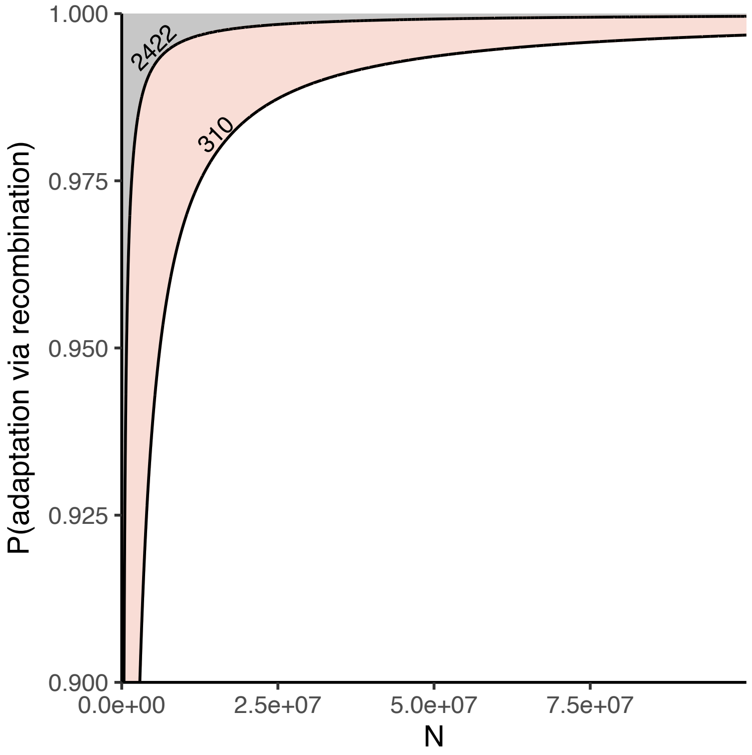

**Figure S11. Adaptations are primarily obtained via recombination in a stochastic model of an across-host sweep.** In the stochastic model, the probability a host adapts from recombination vs. *de novo* mutating an origin is defined in equation 4. We plot here the output of equation 4 in a population of size $10^8$. The pink band corresponds to the range of $r * l_r / \mu$ estimated in common commensal gut bacteria in (Liu and Good 2024), with a lower bound of 310. The grey band denotes all higher values of $r * l_r / \mu$ shown in Fig 4, with a lower bound of 2422. In all ranges, the vast majority of hosts are expected to obtain the adaptation via recombination (> 90% probability).

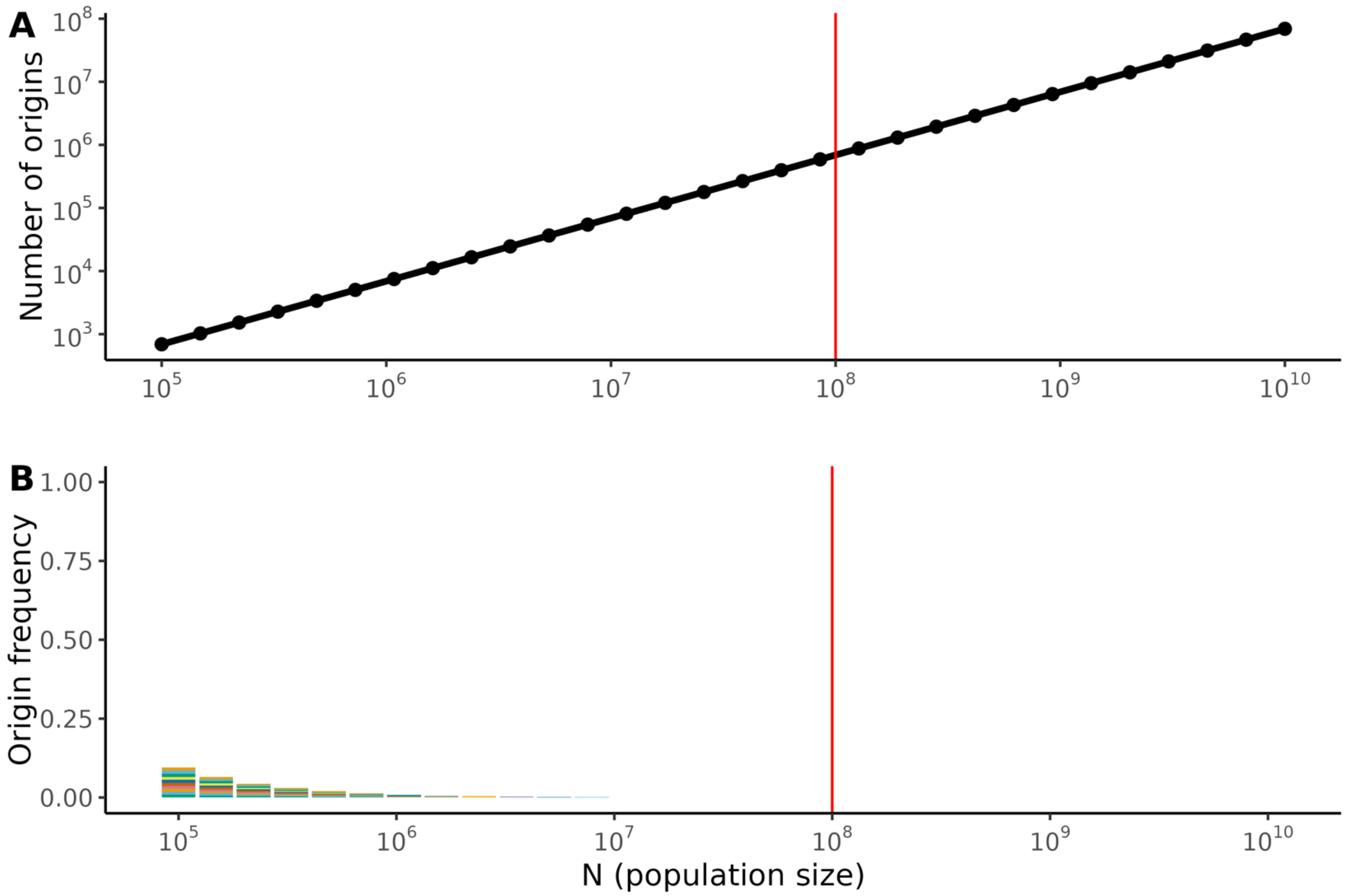

**Figure S12. Number of origins and frequencies of sweeping haplotypes as a function of number of hosts in the population**.

**A.** Number of origins expected in a sweep spreading across hosts and **B.** frequencies of the 10 most frequent haplotypes. We fix $r * l_r / \mu$ at 1000, a value consistent with empirical estimates (e.g. $l_r$ = 10kb, $r / \mu = 0.1$) (Liu and Good 2024). Similar to Fig 4, the number of origins is large, and the frequencies of top origins negligible with a quantity of $r * l_r / \mu$ in the estimated empirical range, even when N varies across five orders of magnitude ($10^5$ - $10^{10}$). We show results in Fig 4 for a population of size $10^8$, denoted here by the red line.

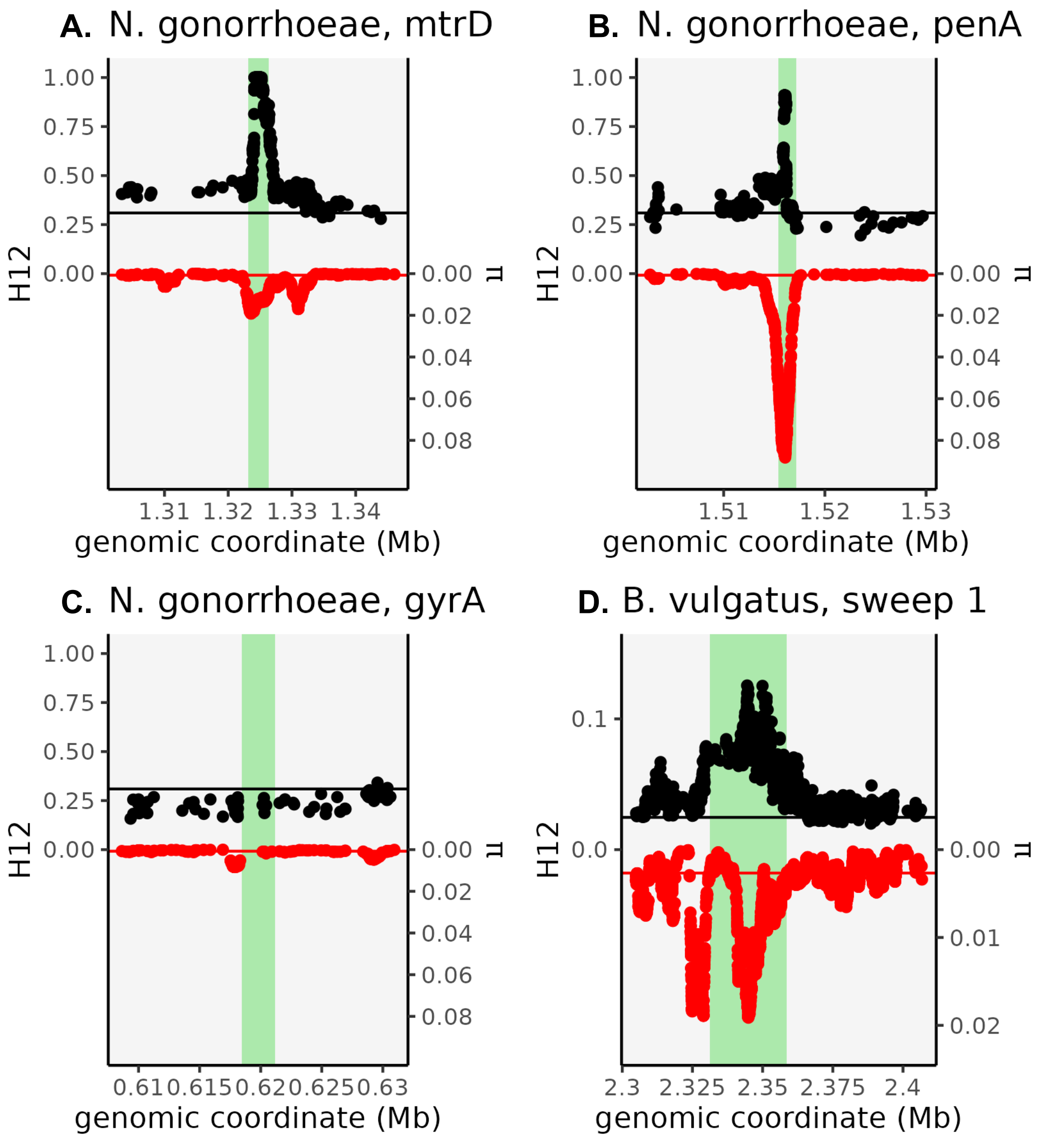

**Figure S13. Local patterns of H12 and nucleotide diversity at three regions in *N. gonorrhoeae* and one sweep in *B. vulgatus*.** Left axis/upper plot denotes H12, while right axis/lower plot denotes π. Both axes are scaled to accommodate the genome-wide maximum value (of H12 or π, respectively). Both the *mtrD* and *penA* genes (green rectangles) in *N. gonorrhoeae* are enriched in π, consistent with their introgressed, mosaic origins (**A-B)**. However, *gyrA* (green rectangle) is enriched neither in π nor H12. (**C**). A sweep in *B. vulgatus* shows a similar enrichment in π and H12, with green rectangle denoting a polysaccharide utilization locus (PUL) in the vicinity (**D**).

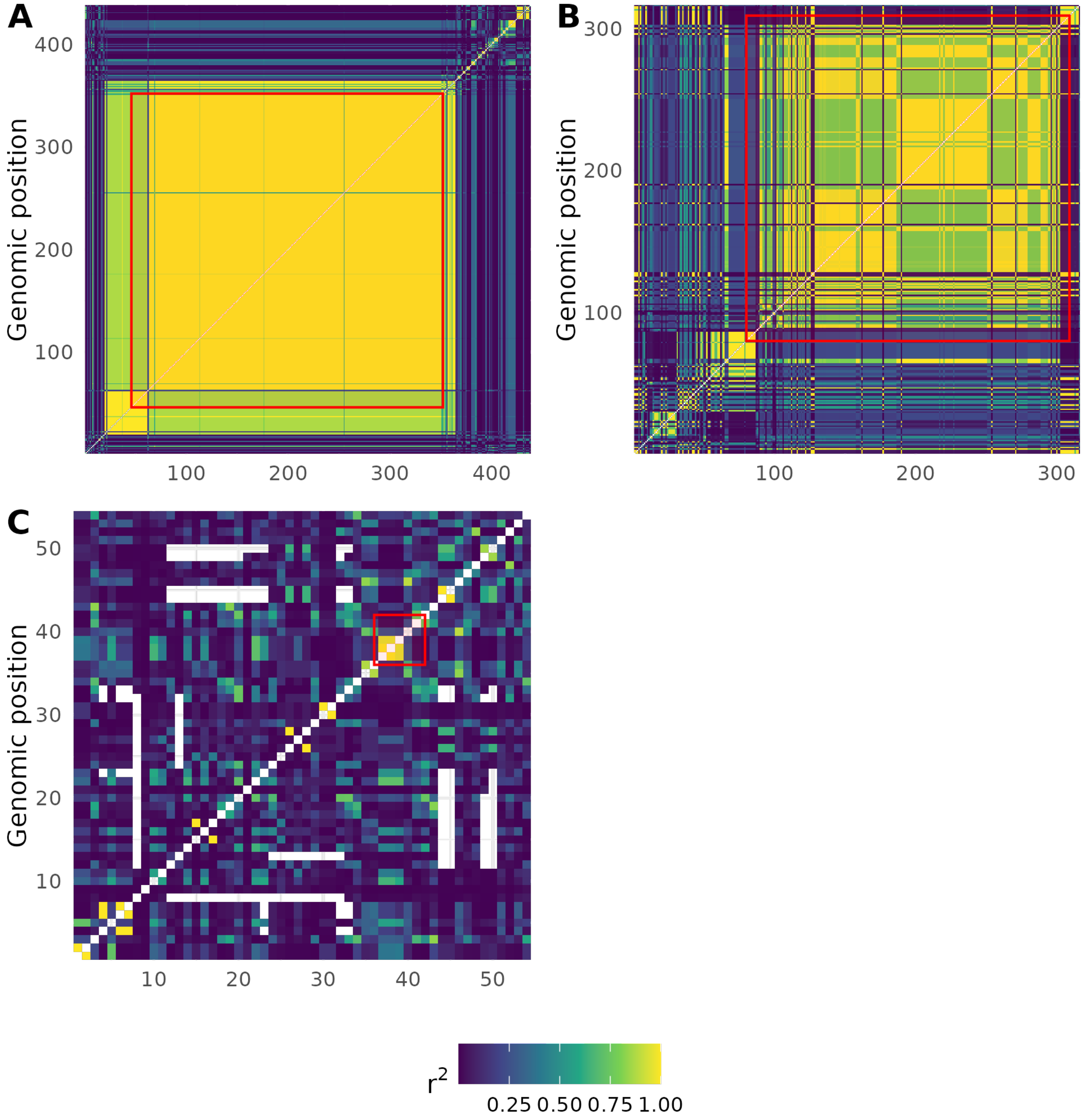

**Figure S14. Local patterns of linkage disequilibrium at *mtr* and *penA* in *N. gonorrhoeae.*** For the *penA*, *mtr,* and *gyrA* regions in *N. gonorrhoeae*, we plot local linkage disequilibrium, requiring variants to have minor allele frequency >= 0.03. **A**. In the *penA* region, near-perfect LD is found over the length of *penA* (red rectangle). **B**. Over *mtr*, LD is highly elevated over the length of the operon (red rectangle). **C.** Over the few variants in *gyrA* (red rectangle), no pervasive elevation in LD is apparent.

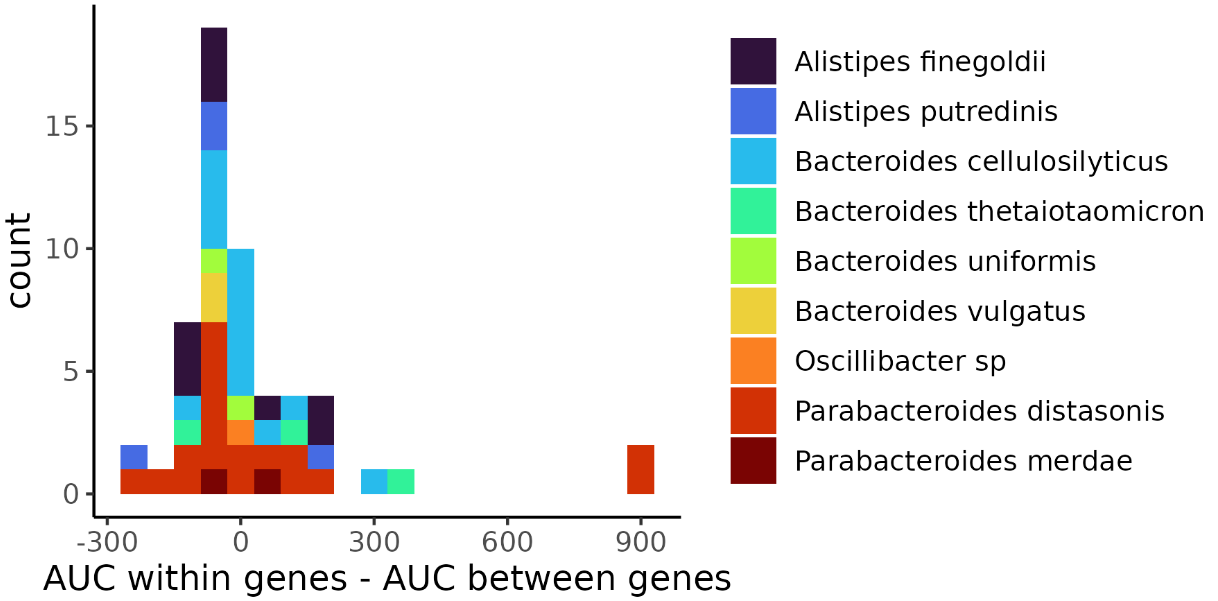

**Figure S15. The difference in linkage disequilibrium AUC within genes vs. between genes for all gut commensal sweeps**

For each sweep, the area under the linkage disequilibrium curve was calculated for pairs of variants residing in the same gene vs. between genes, additionally requiring all variants have minor allele frequency >= 0.03. For sweeps with sufficient comparisons, the difference between the area under the two LD decay curves is shown, with color denoting the species from which the sweep originates. Four sweeps show exceptionally high linkage within genes vs. between (> 2 standard deviations from the mean), with their composite LD decay curve shown in Fig 5D.

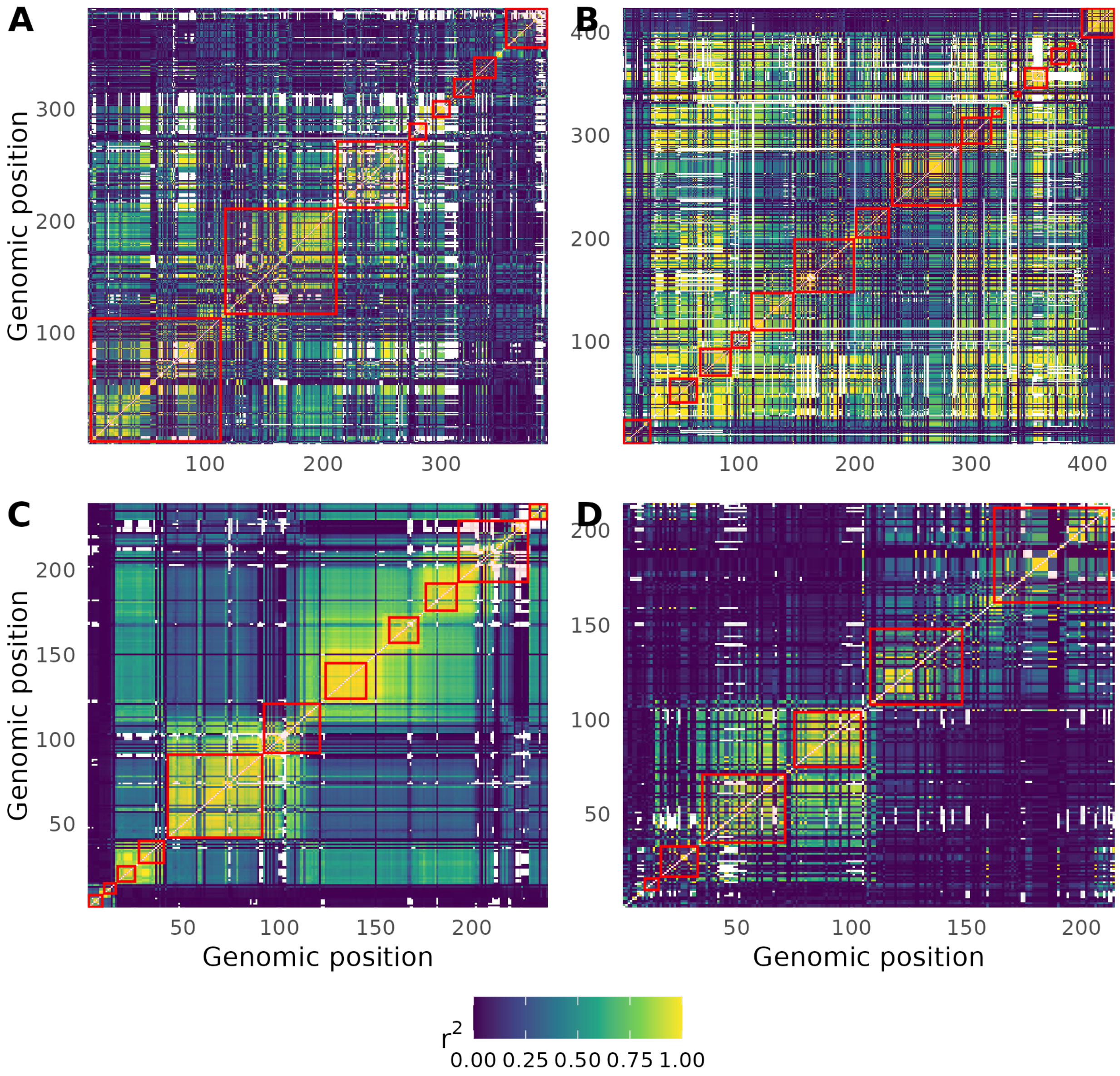

**Figure S16. The maintenance of LD over multiple gene boundaries in four gut commensal sweeps**. For the above sweeps, we highlight regions where high LD seems to be maintained over multiple gene boundaries (denoted with red borders) before decaying precipitously. Such patterns could be consistent with epistasis spanning the region where LD is maintained. (**A**) Sweep in *B. vulgatus* (PUL region), (**B**) sweep in *B. thetaiotamicron*, (**C**). Sweep in *A. putredinis*, (**D**). Sweep in *A. finegoldii*.

#### Supplemental Table Descriptions

**Table S1. H12 Parameters for Selection Scans**
Parameters for the 10 species included in Fig S6. Column 1 corresponds to species name, column 2 to the sample size we downsample to, column 3 to the window length in terms of base pairs and column 4 to the window length as defined in number of SNPs.

**Table S2. Simulation parameters for individual species**
Simulation parameters for the ten species included in Fig S6. Column 1 corresponds to species name. Column 2 corresponds to the tract length estimated from the distribution from Liu and Good Fig 3E (Liu and Good 2024). Column 3 corresponds to the $r / \mu$ value estimated from Liu and Good Fig 1F.

**Table S3. H12 and summary statistics in sweep regions**
Haplotype summary statistics and metadata associated with individual H12 windows in sweep regions. Column 1 corresponds to species name. Column 2 corresponds to the value of H12 for the window. Column 3 corresponds to the contig identity. Column 4 corresponds to the H12 window's midpoint (central SNP) position. Column 5 corresponds to the minimum position of the window. Column 6 corresponds to the maximum position of the window. Column 7 corresponds to the H2/H1 value of the window. Column 8 corresponds to the raw hamming distance between the two most frequent haplotypes, while column 9 corresponds to the hamming12 value (normalized by average genomewide haplotype-divergence). Column 10 corresponds to whether the window is estimated to lie in a genomic region with abnormally low recombination rates (S5 Text). Column 11 corresponds to the gene identity associated with the midpoint position (BV-BRC ID) (Olson et al. 2023). Column 12 corresponds to an arbitrary 'peak' ID (for identifiability across tables). Column 13 corresponds to whether that window is the window with the highest value of H12 for the region (subsequently used for hard / soft inference). Column 14 corresponds to the critical value used to determine significance.

**Table S4. Summary statistics of haplotype structure and BF for sweeps**
Haplotype summary statistics and Bayes factors for each peak classified in Fig 3. Column 1 corresponds to the Bayes factor. Column 2 corresponds to the 'peak' ID. Column 3 corresponds to the H12 value used to classify the peak. Column 4 corresponds to the H2/H1 value used to classify the peak. Column 5 corresponds to the hamming12 value used to classify the peak. Column 6 corresponds to the species name. Column 7 corresponds to the Bayes factor bin used in Fig 3F.

Yee Whye Teh. 2017. "Dirichlet Process." In *Encyclopedia of Machine Learning and Data Mining*.

Zhao, Shijie, Tami D. Lieberman, Mathilde Poyet, et al. 2019. "Adaptive Evolution within Gut Microbiomes of Healthy People." *Cell Host & Microbe* 25 (5): 656-667.e8. https://doi.org/10.1016/J.CHOM.2019.03.007.

Zhou, Siyu, Bo Liu, Dandan Zheng, Lihong Chen, and Jian Yang. 2024. "VFDB 2025: An Integrated Resource for Exploring Anti-Virulence Compounds." *Nucleic Acids Research* 53 (D1): D871–77. https://doi.org/10.1093/nar/gkae968.

Zou, Yuanqiang, Wenbin Xue, Guangwen Luo, et al. 2019. "1,520 Reference Genomes from Cultivated Human Gut Bacteria Enable Functional Microbiome Analyses." *Nature Biotechnology* 37 (2): 179–85. https://doi.org/10.1038/s41587-018-0008-8.
